# The SAGA and NuA4 component Tra1 regulates *Candida albicans* drug resistance and pathogenesis

**DOI:** 10.1101/2021.03.17.435915

**Authors:** Iqra Razzaq, Matthew D. Berg, Yuwei Jiang, Julie Genereaux, Deeva Uthayakumar, Grace H. Kim, Christopher J. Brandl, Patrick Lajoie, Rebecca S. Shapiro

**Affiliations:** Department of Molecular and Cellular Biology, University of Guelph, Guelph, Ontario, Canada N1G2W1; Department of Biochemistry, The University of Western Ontario, London, Ontario, Canada, N6A 5C1; Department of Anatomy and Cell Biology, The University of Western Ontario, London, Ontario, Canada, N6A 5C1

## Abstract

*Candida albicans* is the most common cause of death from fungal infections. Emergence of resistant strains reducing the efficacy of first line therapy with echinocandins such as caspofungin calls for the identification of alternative therapeutic strategies. Tra1 is an essential component of the SAGA and NuA4 transcriptional co-activator complexes. As a PIKK family member, Tra1 is characterized by a C-terminal phosphoinositide 3-kinase domain. In *Saccharomyces cerevisiae,* the assembly and function of SAGA and NuA4 is compromised by a version of Tra1 (Tra1_Q3_) with three arginine residues in the putative ATP-binding cleft changed to glutamine, Whole transcriptome analysis of the *S. cerevisiae tra1_Q3_* strain highlights Tra1’s role in global transcription, stress response and cell wall integrity. As a result, *tra1_Q3_* increases susceptibility to multiple stressors, including caspofungin. Moreover, the same *tra1_Q3_* allele in the pathogenic yeast *Candida albicans* causes similar phenotypes, suggesting that Tra1 broadly mediates the antifungal response across yeast species. Transcriptional profiling in *C. albicans* identified 68 genes that were differentially expressed when the *tra1_Q3_* strain was treated with caspofungin, as compared to gene expression changes induced by either *tra1_Q3_* or caspofungin alone. Included in this set were genes involved in cell wall maintenance, adhesion and filamentous growth. Indeed, the *tra1_Q3_* allele reduces filamentation and other pathogenesis traits in *C. albicans*. We identified *EVP1*, which encodes a putative plasma membrane protein, amongst the Tra1-regulated genes, Disrupting *EVP1* results in reduced filamentation and infection capacity in *C. albicans*. Thus,Tra1 emerges as a promising therapeutic target for fungal infections.

**Importance:** Fungal pathogens such as *Candida albicans* are important agents of infectious disease, with increasing rates of drug resistance, and limited available antifungal therapeutics. In this study, we characterize the role of *C. albicans* Tra1, a critical component of acetyltransferase complexes, involved in transcriptional regulation and responses to environmental stress. We find *C. albicans* genetic mutants with impaired Tra1 function have reduced tolerance to cell-wall targeting stressors, including the clinically-important antifungal caspofungin. We further use RNA-sequencing to profile the global fungal response to the *tra1* mutation, and identify a previously uncharacterized *C. albicans* gene, *EVP1*. We find that both *TRA1* and *EVP1* play an important role in phenotypes associated with fungal pathogenesis, including cellular morphogenesis, biofilm formation, and toxicity towards host immune cells. Together, this work describes the key role for Tra1 in regulating fungal drug tolerance and pathogenesis, and positions this protein as a promising therapeutic target for fungal infections.

## Introduction

Fungal infections are a major modern public health challenge, killing over one million people annually (1, 2). While the human immune system presents an effective barrier to infection in healthy individuals, most fungal pathogens are opportunistic and can cause deadly invasive infections in immunosuppressed patients. *Candida* species are amongst the most common causes of life-threatening invasive fungal infections, with *Candida albicans* being the most frequently isolated pathogen (3, 4). Currently, there is a limited armamentarium of effective, non-toxic antifungal therapeutics for the treatment of invasive candidiasis, with three major classes of antifungal drugs in clinical use: polyenes, azoles, and echinocandins (5). However, acquired resistance to these antifungal agents is increasingly common amongst *C. albicans* isolates, and non-*albicans Candida* species with high rates of acquired or intrinsic antifungal resistance (6, 7), including the emerging multidrug-resistant pathogen *Candida auris*, becoming increasingly prevalent (8–11).

The echinocandins, which include caspofungin, are the most recently discovered class of antifungal drugs and are fungicidal against most *Candida* pathogens (12). Echinocandins cause significant cell wall stress by inhibiting the (1, 3)-β-d-glucan synthase (encoded in *Candida* by *FKS1/2),* and thus decrease production of the critical cell wall component (1, 3)-β-d-glucan (13, 14). In many cases, echinocandins are the first line of treatment for invasive candidiasis, and the majority of patients with candidemia receive echinocandins (15–17). As a result, a growing proportion of disease-causing *Candida* isolates display reduced susceptibility to echinocandins, and in some cases, cross-resistance with other antifungals such as azoles (16, 18, 19). The increasing threat of antifungal resistance, in combination with the already limited diversity of antifungal drug classes, make it imperative to discover alternative lines of treatment and expand the availability of effective therapeutic approaches.

Antifungal drugs such as the echinocandins activate fungal stress responses, which are important in the susceptibility and resistance to antifungal therapeutics (20–23). As with other stress responses, these antifungal stress response pathways are controlled through gene expression. In eukaryotic organisms, chromatin modifications are critical for the regulation of gene expression (24, 25). One role of modifications, such as the acetylation of lysine residues on the N-terminal tails of histones, is to facilitate the recruitment of the transcriptional machinery to target promoters. The SAGA (Spt-Ada-Gcn5-Acetyltransferase) and NuA4 (Nucleosome acetyltransferase of H4) complexes contain two of the principal lysine acetyltransferases, Gcn5 and Esa1, respectively (26–28). Both target lysines within the N-terminal tails of histones and have substrates with roles unrelated to chromatin (29–31). The SAGA complex has a broad role in transcription, including regulating multiple stress response pathways (32–34).

Tra1/TRRAP is an essential component of SAGA and NuA4 complexes (35–37). It is a member of the PIKK (phosphoinositide-3-kinase-related kinase) family that also includes Tor1, ATM and Rad3-related protein (ATR/Mec1), and the DNA-dependent protein kinase catalytic subunit (DNA-PKcs) (38). As with other PIKKs, Tra1 is incorporated into SAGA and NuA4 in a process that requires Hsp90 and its co-chaperone, the Triple-T complex (TTT complex; (39, 40). Tra1 contains four domains: an N-terminal HEAT region, followed by FAT, PI3K, and FATC domains (38, 41–43). The PI3K domain is essential for Tra1 function (43, 44), yet unlike other PIKKs (45), Tra1 lacks kinase activity (36, 46). We recently identified three arginine residues proximal to what is the ATP-binding cleft in the PI3K domain of other PIKK proteins, that, when mutated to glutamine, reduce growth, impair assembly of SAGA and NuA4 complex, and cause dysregulated transcription; we termed this allele *tra1_Q3_* (47). In the model yeast *Saccharomyces cerevisiae*, the *tra1_Q3_* allele results in sensitivity to high temperature, inositol auxotrophy, and decreased resistance to cell wall perturbations (47). The arginine residues are conserved among Tra1 orthologues, including in other yeast species such as *C. albicans* (47).

Given the connection between transcriptional regulation and stress response, it follows that chromatin-modifying factors would have important functions in mediating fungal pathogenesis, including their virulence and susceptibility to antifungal drugs (48). Indeed, chromatin-modifying factors including lysine (K) acetyl-transferases (KATs) such as Gcn5, Hat1, and Rtt109, and lysine deacetylases (KDACs) such as Set3 and Rpd31 are implicated in *Candida* virulence processes including filamentous growth and biofilm formation (49–56), as well as in resistance and susceptibility to antifungal drugs (52, 57–61). The requirement of Tra1 for resistance to cell wall perturbations in *S. cerevisiae* (47), along with its role in both SAGA and NuA4 complexes, positions Tra1 as a potentially unique target for antifungal therapy. Despite this, the ortholog of *TRA1* in the fungal pathogen *C. albicans* remains to be characterized and its role in antifungal resistance and fungal virulence has not been explored.

Here, we identify a key role for Tra1 in mediating resistance to antifungal drugs, as well as fungal virulence. We demonstrate that the *tra1_Q3_* allele increases sensitivity to the antifungal echinocandin caspofungin in *S. cerevisiae*. Using CRISPR editing to generate an orthologous *tra1_Q3_* mutant in *C. albicans* and a CRISPR interference (CRISPRi) based *tra1* repression strain, we show that diminishing Tra1 increases sensitivity to caspofungin and other cell wall stressors in this fungal pathogen. Transcriptome profiling in *C. albicans* revealed that Tra1 is required for the transcriptional changes that occur in response to caspofungin treatment. Many of these genes were also involved in *C. albicans* morphogenesis and pathogenesis. We find that expression of C3_07470W (*EVP1*) — a previously uncharacterized *C. albicans* gene — was significantly increased in the *tra1_Q3_* strain in the presence of caspofungin. As is the case with diminished Tra1 function, disrupting *EVP1* results in a *C. albicans* strain with reduced filamentation capacity and infection potential in a macrophage infection model. Together, this work positions Tra1 as a key regulator of *C. albicans* drug resistance and pathogenesis, and highlights its potential as a target for antifungal therapeutics.

## Results

### TRA1 mutations increase susceptibility to caspofungin in S. cerevisiae and C. albicans

Amongst Tra1 orthologues, the positively charged residues at positions 3389, 3390 and 3456 are highly conserved (47). When mapped onto the cryo-EM structure of Tra1 (42), these conserved arginine residues fall within or near the ATP-binding cleft (47). Although the PI3K domain of Tra1 lacks key residues required for ATP binding, altering these three arginine residues to glutamine, an allele termed *tra1_Q3_*, reduces assembly of Tra1 into SAGA and NuA4 complexes in *S. cerevisiae,* demonstrating the importance of the cleft (47). To further assess the importance of the PI3K domain of Tra1 on SAGA/NuA4-mediated transcription, we performed RNA sequencing (RNA-seq) to identify genes and pathways that are differentially expressed in the *S. cerevisiae tra1_Q3_* strain compared to a wild-type strain. We identified 1724 and 1792 genes that were significantly up- and downregulated, respectively (adjusted *P*-value < 0.05; **FIGURE 1a and SUPP FILE 1**). In the set of genes upregulated greater than 2-fold in *tra1_Q3_* relative to wild-type, GO term analysis identified enrichment for genes involved in cell wall organization (**FIGURE 1b)**. Down regulated genes with greater than 2-fold change were enriched for roles in polyphosphate metabolism (**FIGURE 1c)**.

**Figure 1.**
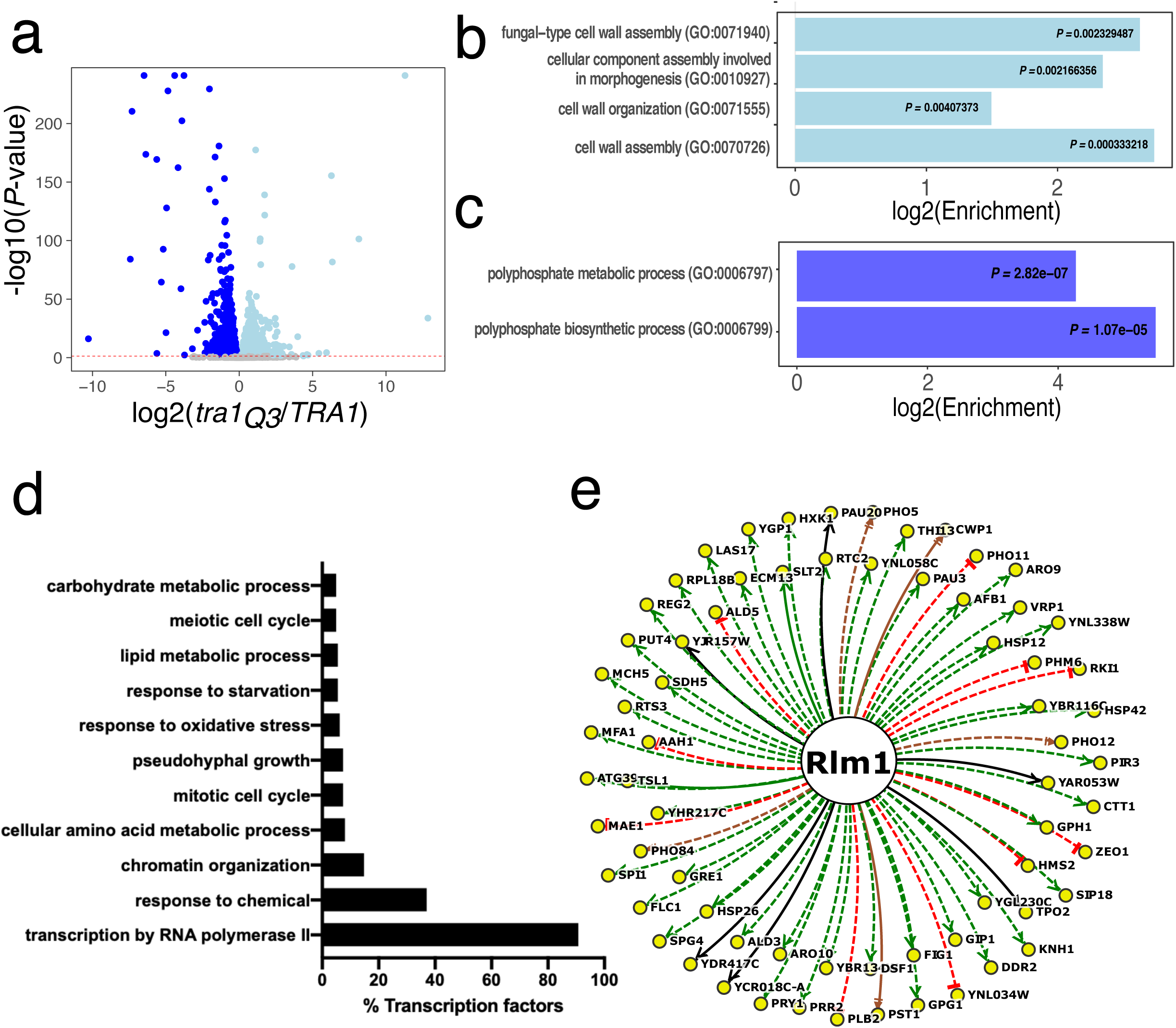
Transcriptome analysis of *S. cerevisiae* expressing *tra1_Q3_*. **(a)** Volcano plot of changes in gene expression as determined by RNA-seq in strains expressing *tra1_Q3_* relative to a strain expressing wild type *TRA1*. Differentially expressed genes with an adjusted *P*-value < 0.05 are colored light blue (up regulated in *tra1_Q3_* relative to *TRA1*) and dark blue (down regulated in *tra1_Q3_* relative to *TRA1)*. **(b)** Significantly enriched GO biological processes were determined from the set of significant genes with > 2-fold up regulation (adjusted *P* < 0.05) in *tra1_Q3_* relative to *TRA1*. **(c)** Significantly enriched GO biological processes were determined from the set of genes with > 2-fold down regulation (adjusted *P* < 0.05) in *tra1_Q3_* relative to *TRA1*. **(d)** Differentially expressed genes in *tra1_Q3_* were analysed for transcription factor associations using YEASTRACT. GO analysis was performed with the percentage of identified transcription factors (162) for each GO term shown in the bar graph. **(e)** Rlm1 associations with differentially expressed genes in the *tra1_Q3_* strain. The experimental evidence underlying each regulatory association (solid lines for DNA-binding evidence; dashed lines for expression evidence), as well as the sign of the interaction—positive (green), negative (red), positive and negative (brown), or undefined (black) are displayed.

We also identified differential expression of genes involved in various stress responses in the *tra1_Q3_* strain, in agreement with previous studies demonstrating that SAGA and NuA4 regulate stress responsive genes (32, 62, 63). These included genes involved in the cell wall integrity pathway (64) (*PKH1, MTL1, KSS1, SWI6, MSG5, WSC3, PTP2* and *PTP3* were downregulated and *SAC7* and *RHO1* were upregulated). Previous work also demonstrated a role for Tra1 in resistance to cell wall stress (44, 47, 65, 66). We then analyzed differentially regulated genes (392 genes, log_2_ fold change > 1, *P* < 0.05) for known association with transcription factors using yeastract (www.yeastract.org)(67) (**FIGURE 1d**, **SUPP FILE 2**). GO analysis of identified transcription factors revealed factors involved in PolII transcription, in agreement with the broad role of SAGA in global transcriptional regulation (34). Other transcription factor associations (Hsf1, Hac1) highlight the role of *TRA1* in stress responses. Interestingly, our analysis also shows significant enrichment in Sfp1 target genes and *RLM1* targets (**FIGURE 1e**). Sfp1 interacts with Tra1 (68) and we recently demonstrated that Sfp1 regulates Tra1/SAGA-driven gene expression in response to protein misfolding (69). *RLM1* controls gene expression of factors involved in the cell wall integrity pathway (CWI) (64), consistent with the *tra1_Q3_* causing increased sensitivity to cell wall perturbation (47). Moreover, SAGA recruitment to the promoters of CWI target genes requires *RLM1*. (65).

As many antifungals target the cell wall or plasma membrane and induce stress responses (70), we hypothesized that *TRA1* might regulate the antifungal response. We therefore assessed whether the *tra1_Q3_* strain was sensitive to the antifungal agent caspofungin, an echinocandin that causes cell wall stress. Growth of the *tra1_Q3_* strain was significantly reduced by caspofungin in both solid and liquid media conditions **(FIGURE 2a, b)**.

**Figure 2.**
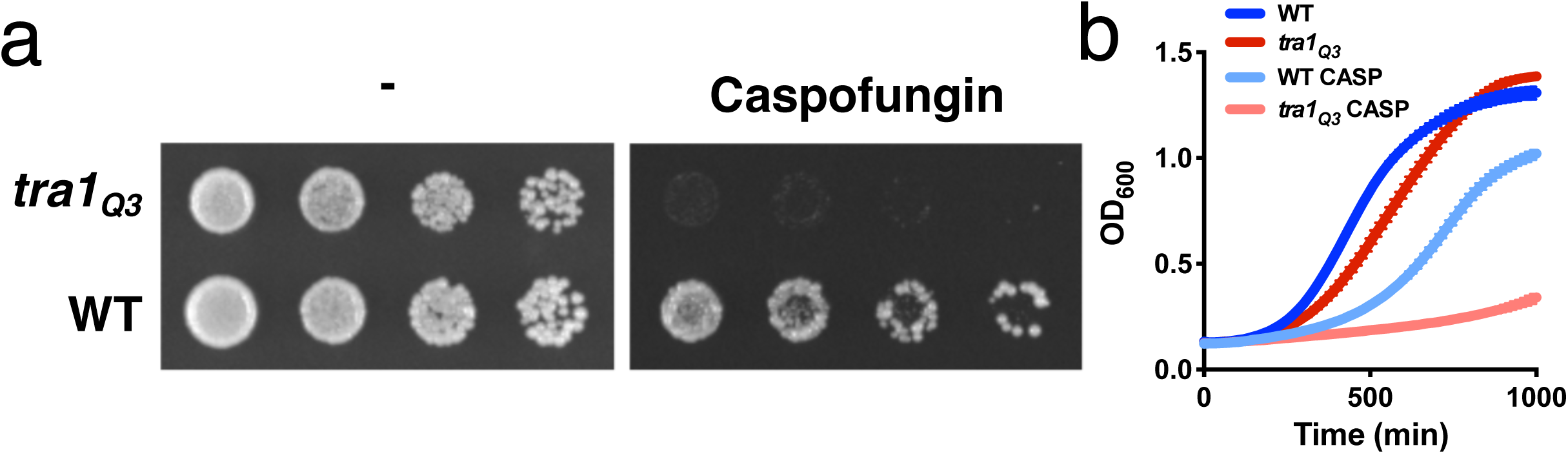
*S. cerevisiae TRA1* deficient strains are hypersensitive to cell wall stressors. **(a)** Growth of the *S. cerevisiae tra1_Q3_* strain on solid medium is impaired by 0.25 µg/mL caspofungin compared to a wild-type strain. *S. cerevisiae* strains were serially diluted (1:10) and spotted onto solid YPD agar media with or without 0.25 µg/mL caspofungin. **(b)** Growth of the *S. cerevisiae tra1_Q3_* strain is hypersensitive to 0.25 µg/mL caspofungin in liquid medium. *S. cerevisiae* strains were grown in liquid YPD medium with or without 0.25 µg/mL caspofungin continuously over 1000 minutes. Growth was determined by optical density (OD_600_).

Hypothesizing that *TRA1* could be a potential target for antifungal therapy, we next examined whether *TRA1* was similarly involved in sensitivity to cell wall stressors in the fungal pathogen *C. albicans*. First, we identified the three conserved positively charged residues (K3471, R3472 and R3538) that correspond to the residues mutated in the *S. cerevisiae tra1_Q3_* (**FIGURE 3a**). We used CRISPR-Cas9-based genome editing tools (71) to modify these residues to glutamine for both alleles of *TRA1* in diploid *C. albicans*. We generated two independent *tra1_Q3_* strains in *C. albicans* and assessed the sensitivity of these strains to cell wall stressors, as compared with a wild-type strain. As was the case in *S. cerevisiae*, *tra1_Q3_* rendered *C. albicans* significantly more sensitive to the presence of the cell wall stressor calcofluor white, as well as the cell wall-targeting antifungal caspofungin (**FIGURE 3b**).

**Figure 3.**
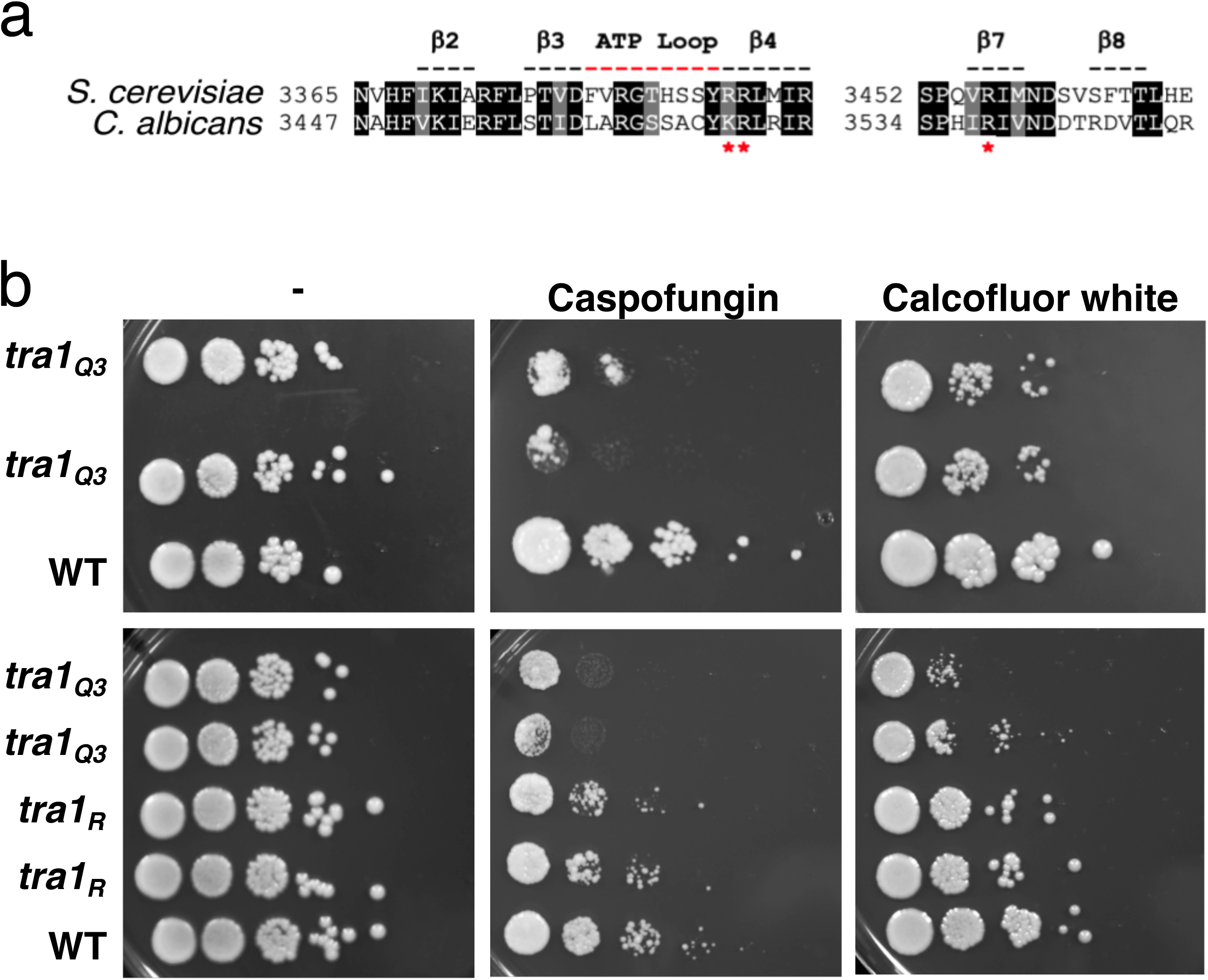
*C. albicans tra1* strains are hypersensitive to cell wall stressors. **(a)** Alignment of *TRA1* from *S. cerevisiae* and *C. albicans*, highlighting (red asterisks) the conserved positively charged residues (K3471, R3472 and R3538) proximal to the PI3K ATP-binding cleft, which correspond to residues mutated in the *S. cerevisiae tra1_Q3_* strain. **(b)** *tra1_Q3_* results in hypersensitivity caspofungin or calcofluor white in solid medium. *C. albicans* strains were serially diluted (1:10 dilutions) and spotted onto solid YPD agar media with or without 0.25 µg/mL caspofungin or 25 µg/mL calcofluor white. **(c)** Growth of *C. albicans tra1* CRISPRi repression strains (“*tra1_R_*”) on solid medium containing 0.25 µg/mL caspofungin or 25 µg/mL calcofluor white. *C. albicans* strains were serially diluted (1:10 dilutions) and spotted onto solid YPD agar media with or without either drug.

To confirm the role of *TRA1* in susceptibility to cell wall stress, we generated a *TRA1* genetic depletion strain using a CRISPR interference (CRISPRi)-based repression system optimized for *C. albicans* (72). We monitored sensitivity of the CRISPRi *tra1* repression strains (*tra1_R_*) compared with a wild-type strain, and found that *TRA1* repression increased sensitivity to calcofluor white and caspofungin, though less so than the *tra1_Q3_* allele (**FIGURE 3c**). Together, these results confirm the importance of Tra1 in protecting the fungal cell from cell wall-associated stress, including the antifungal caspofungin.

### RNA-seq analysis reveals the transcriptional signature of tra1_Q3_ in response to caspofungin

Given the role of Tra1 as an essential component of SAGA and NuA4 histone acetyltransferase complexes, and its role in sensitivity to the antifungal caspofungin, we next sought to explore the role of *TRA1* in mediating the fungal transcriptional response to treatment with caspofungin. Wild-type and *tra1_Q3_* strains of *C. albicans* were grown in the presence or absence of caspofungin, and RNA-seq was performed. Overall, ∼45% of the variation in the resultant transcriptomic dataset was explained by treatment with caspofungin, and ∼20% of the variation explained by the *tra1_Q3_* mutation (**FIGURE 4a**). For the untreated *tra1_Q3_* strain, 1514 genes were differentially expressed, 841 downregulated and 673 upregulated when compared to wild-type (adjusted *P*-value < 0.05). Caspofungin treatment in wild-type cells resulted in 3113 genes being differentially expressed, 1723 downregulated and 1390 upregulated. Compared to the wild-type *C. albicans* strain, the number of genes differentially regulated upon caspofungin treatment in the *tra1_Q3_* expressing strain was reduced to 1763, 766 downregulated and 997 upregulated. A comprehensive list of significant genes and the fold-change expression data for each comparison can be found in **SUPP FILE 3**.

**Figure 4.**
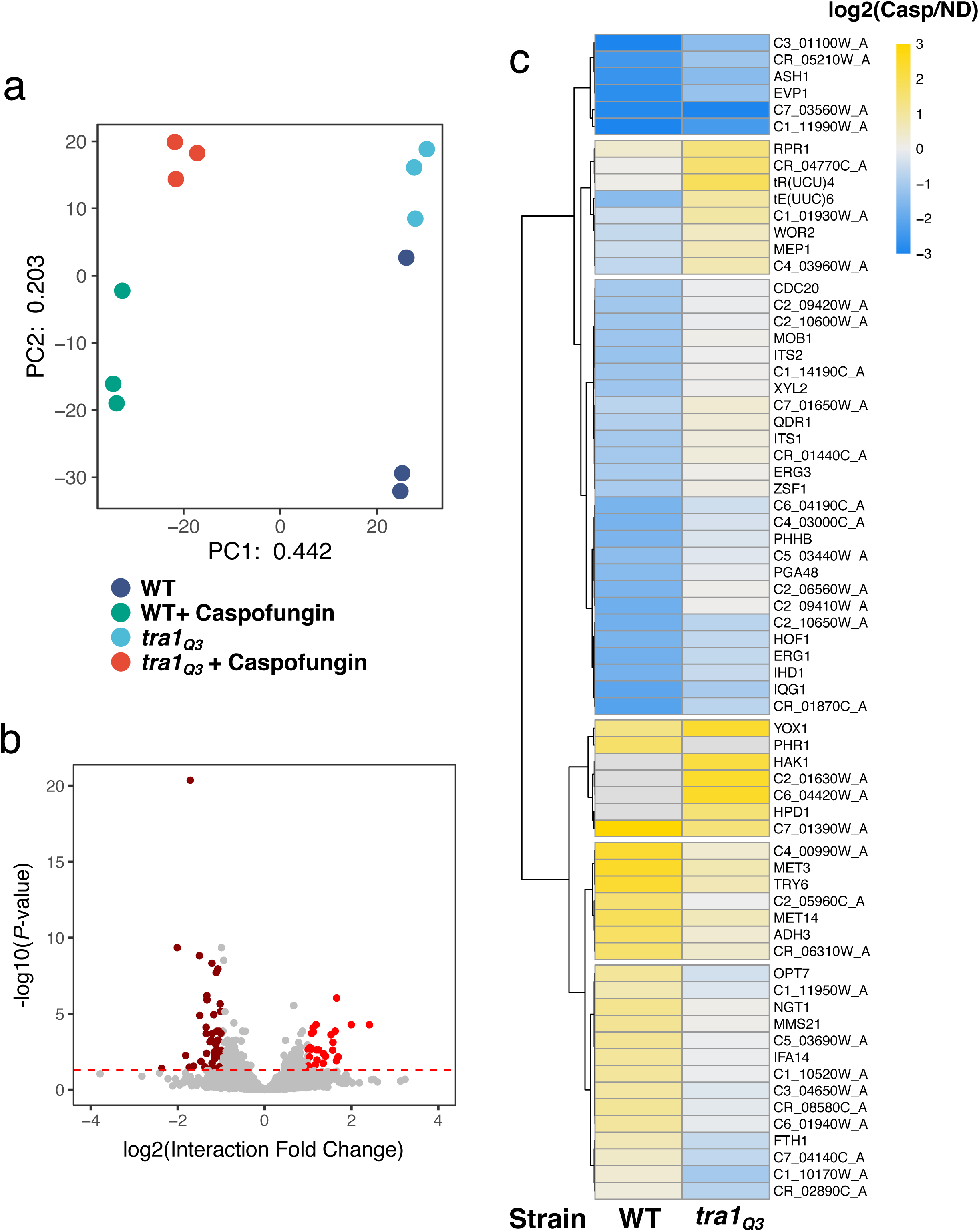
Transcriptome analysis of *C. albicans* expressing *tra1_Q3_* in response to caspofungin. **(a)** Principal component analysis of centered log ratio normalized reads from *C. albican* expressing either wild type *TRA1* or *tra1_Q3_* with and without caspofungin treatment. Each point represents one biological replicate (n = 3). **(b)** Volcano plot of genes that respond differently to caspofungin treatment in the strain expressing *tra1_Q3_* relative to the wild type strain. Genes with log_2_ fold change response > 1 (adjusted *P*-value < 0.05) are colored red and genes with log_2_ fold change response < -1 (adjusted *P*-value < 0.05) are colored dark red. **(c)** Heatmap of hierarchical clustered genes with different responses to caspofungin in *tra1_Q3_* expressing strains relative to wild type strains (log_2_ fold change > |1|, adjusted *P*-value < 0.05). Fold-change for each gene is the average from three biological replicates. Genes that are upregulated and downregulated upon caspofungin treatment are colored yellow and blue, respectively.

From this dataset, we identified the genes responding most differently to treatment with caspofungin in the presence of *tra1_Q3_*, as compared to caspofungin treatment of the wild-type strain and thus possibly underpinning the genotype-condition interaction (**FIGURE 4b**). There were 68 genes that had a significantly different response based on this genotype-condition interaction (log_2_ fold change > 1, adjusted *P*-value < 0.05). A heat map representing the changes in expression upon caspofungin treatment for wild type and *tra1_Q3_* strains is shown in **FIGURE 4c**. The most pronounced differences were seen for *HPD1*, a dehydrogenase involved in the degradation of toxic propionyl-CoA, and *PHR1,* a cell surface glycosidase. Although there was no significant Gene Ontology (GO) term enrichment in this smaller subset of differentially regulated genes, the genes annotated with a known biological process included many involved in cellular transport (*ADH3, EVP1, FTH1, HAK1, HOF1, HPD1, MEP1, MOB1, NGT1, OPT7, PHHB, QDR1, TRY6*), filamentous growth (*ASH1, ERG1, ERG3, PHHB, PHR1, WOR2*), biofilm growth (*PHR1, QDR1, TRY6*), and response to chemicals, drugs, and stress (*ASH1, ERG1, ERG3, HOF1, MEP1, MMS21, PHHB, HAK1, PHR1, QDR1*).

The genes uniquely regulated by *TRA1* upon caspofungin treatment were then analyzed for regulatory associations (including documented DNA-binding and expression-based evidence) with known transcription factors using the PathoYeastract portal (67). We identified several transcription factors including Pho4 (known for being regulated by SAGA/NuA4 in *S. cerevisiae* (*73–75*)) as well as Nrg1 and Ron1, with known roles in filamentous growth, biofilm formation and response to stress and hyphal growth (**FIGURE 5a and SUPP FILE 4**)(76–80). Regulatory associations of Pho4, Nrg1 and Ron1 with genes differentially expressed in *tra1_Q3_* are shown in **FIGURE 5b**. Other transcription factors previously linked to SAGA/NuA4 include Snf5, Skn7, Sko1, Rim101, Fkh2 and Mcm1 (60, 81–84). Among identified factors, Sko1 and Cas5, have been shown to regulate the response to the caspofungin-induced cell wall damage in *C. albicans* (*85–87*). Taken together, these data support an important role for Tra1 in the regulation of the transcriptional response to caspofungin.

**Figure 5.**
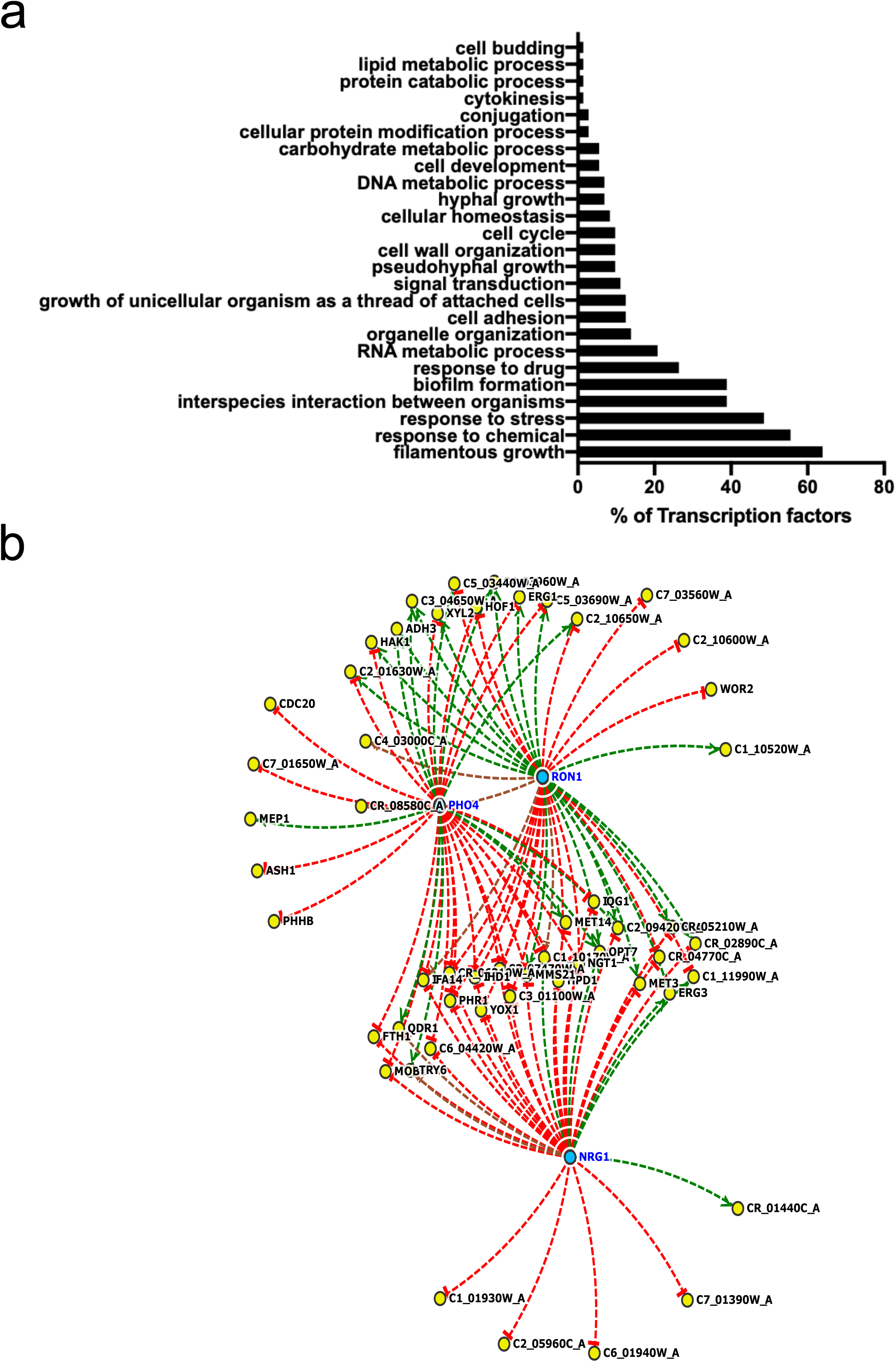
Transcription factor association underpinning the *tra1_Q3_* response to caspofungin in *C. albicans*. **(a)** Differentially expressed genes in *tra1_Q3_* in response to caspofungin treatment were analysed for transcription factor associations using YEASTRACT. GO analysis was performed and the percentage of identified transcription factors (73) for each GO term shown in the bar graph. **(b)** Pho4, Nrg1 and Ron1 associations with differentially expressed genes in the *tra1_Q3_* strain in response to caspofungin treatment. The experimental evidence underlying each regulatory association (full lines for DNA-binding evidence; dashed for expression evidence), as well as the sign of the interaction—positive (green), negative (red), positive and negative (brown) or undefined (black) are displayed.

### TRA1 plays a role in C. albicans pathogenesis

Since many differentially regulated genes unique to the *tra1_Q3_* response to caspofungin had annotated roles in fungal pathogenesis processes, we explored whether *TRA1* is involved in pathogenesis phenotypes in *C. albicans*. One key pathogenesis-associated phenotype for *C. albicans* is the ability to undergo a reversible morphological transition between yeast and filamentous growth (70). We monitored wild-type and *tra1_Q3_* strains of *C. albicans* for their ability to undergo cellular morphogenesis and filamentation in response to diverse media conditions known to induce filamentation, including growth in medium containing 10% serum and Spider medium. We found that filamentation of the *tra1_Q3_* strain isolates was impaired compared to the wild-type strain (**FIGURE 6a**), suggesting that *TRA1* is required for the filamentous growth transition in *C. albicans*. This filamentous growth defect is specific to filamentation, and is not an overall growth defect of this mutant (**FIGURE S1**). Given the important role that morphological transitions play in biofilm formation in *C. albicans* (88), we further investigated the role of *TRA1* in biofilm formation. We allowed *C. albicans* biofilms to form in multi-well plates, removed planktonic cells, and quantified the presence of metabolically-active adherent fungal cells as a measure of biofilm growth. The *tra1_Q3_* isolates were less capable of forming robust biofilms, compared to a wild-type strain of *C. albicans* (**FIGURE 6b**; *P <* 0.0001, ANOVA).

**Figure 6.**
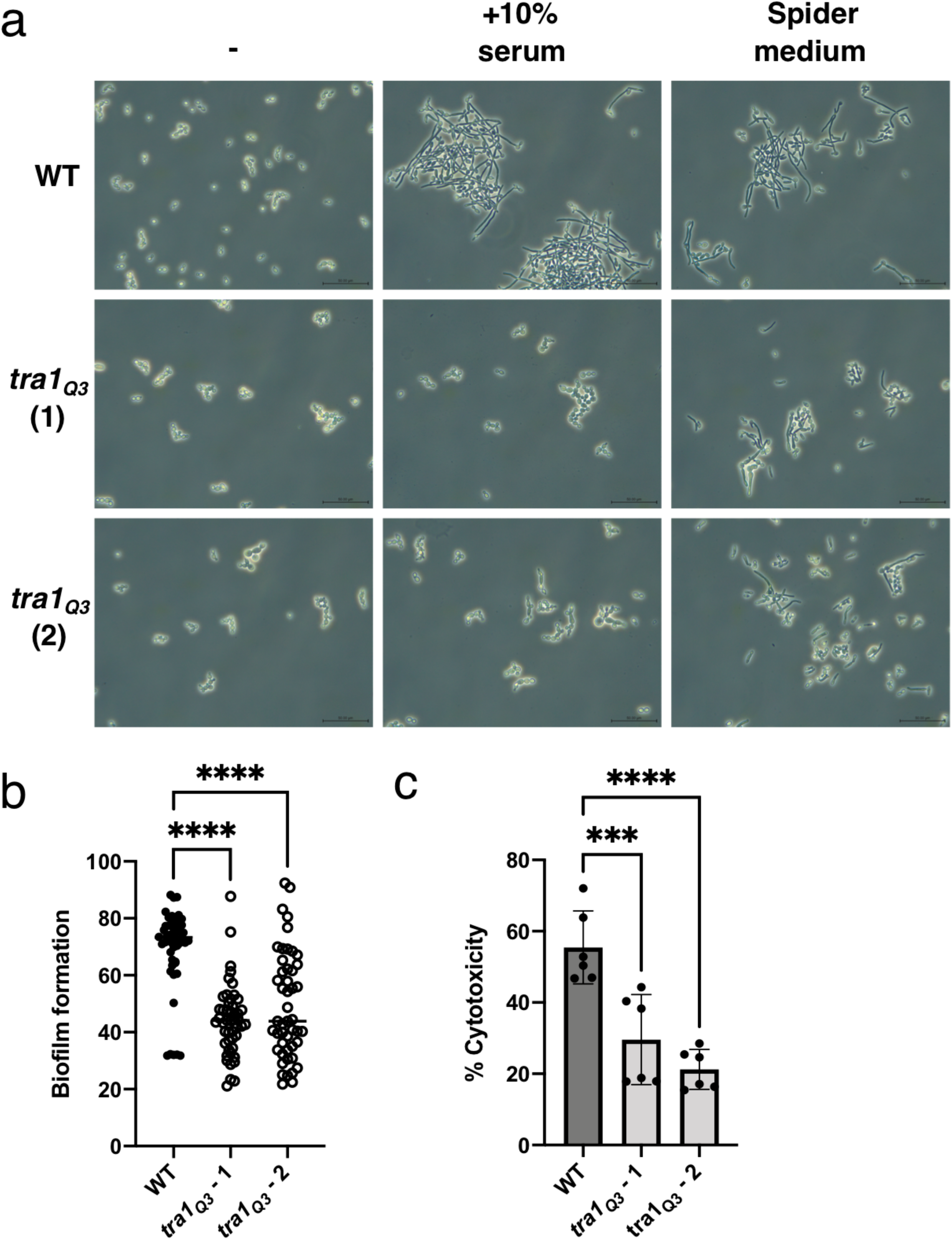
In *C. albicans tra1_Q3_* impairs pathogenesis-associated traits. **(a)** *C. albicans tra1_Q3_* mutants are impaired in the morphogenetic transition from yeast to filamentous growth in YPD media containing 10% serum or in Spider media. *C. albicans* strains were grown in YPD media alone at 30°C, or in YPD containing 10% serum or in Spider media at 37°C for 4 hours. Cells were observed by bright field microscopy. **(b)** *C. albicans tra1_Q3_* mutants are impaired in biofilm formation relative to a wild-type strain (ANOVA, P < 0.0001 (****)). Biofilm growth was quantified by an XTT metabolic readout, measured at OD_490_ and normalized to planktonic growth. Biofilm formation of these fungal strains were measured alongside the *evp1Δ* mutant in Figure 5, and thus the wild-strain measurements are the same between these two figures. **(c)** The *C. albicans tra1_Q3_* strain has reduced macrophage cytotoxicity relative to a wild-type strain (ANOVA, *P* < 0.0001 (****), *P* < 0.001 (***), error bars represent standard deviation). Macrophage cytotoxicity was quantified by measuring LDH after 6 hours of co-incubation with *C. albicans* strains. For LDH measurement, OD_490_ was measured, and the percent cytotoxicity of each strain calculated using these OD_490_ values relative to OD_490_ of uninfected control macrophages. Cytotoxicity associated with these fungal strains was measured alongside the *evp1Δ* mutant in Figure 7, and thus the wild-strain measurements are the same between these two figures.

The importance of *TRA1* in mediating filamentation and in biofilm formation, led us to determine whether it was also involved in the interaction of *C. albicans* with mammalian host cells, as *C. albicans* morphogenesis is linked with host cell interaction and escape (89). We performed a macrophage-*C. albicans* infection assay, where fungal cells were co-incubated with BALB/c immortalized murine macrophage cells. We monitored lactate dehydrogenase (LDH) release as a measure of macrophage cell death upon infection by wild-type or *tra1_Q3_ C. albicans* strains, relative to uninfected macrophages. Macrophage cells infected with *tra1_Q3_* strains generated significantly less LDH compared to macrophages infected with wild-type *C. albicans* (**FIGURE 6c**; *P <* 0.001, ANOVA), indicating less host cytotoxicity associated with *tra1_Q3_*, and suggesting a role for *TRA1* in the infection and killing of host macrophage cells.

### Characterizing EVP1 as a novel regulator of fungal pathogenesis and drug resistance

We next asked whether the transcriptomic data might identify other novel regulators of pathogenesis and antifungal drug resistance. Many of the 68 caspofungin-responsive genes modulated 2-fold or more by *TRA1* are uncharacterized. We focused on C3_07470W (*EVP1*), an uncharacterized *C. albicans* gene encoding a putative plasma membrane protein with a predicted role in cell wall integrity (90). *EVP1* was slightly downregulated in the untreated *tra1_Q3_* strain compared to the untreated wild-type strain and significantly upregulated in the *tra1_Q3_* mutant compared to the wild-type strains when treated with caspofungin (**FIGURE S2**). An *EVP1* mutant *C. albicans* strain has not previously been generated, but Dawon *et al*. (91) recently detected the protein in high abundance in extracellular vesicle (EV) samples, and designated it ‘EV-associated Protein 1ʹ or Evp1.

To study the phenotypes associated with this uncharacterized gene, we used a CRISPR deletion construct to delete *EVP1* in *C. albicans*. We first assessed whether the *evp1Δ/evp1Δ* strain (‘*evp1Δ’*) had altered resistance to cell wall stressors. Wild-type, *evp1Δ* and *tra1_Q3_* strains were grown on medium containing calcofluor white or caspofungin. Unlike the *tra1_Q3_* strain, which is highly sensitive to these stressors, the sensitivity of the *evp1Δ* strain to either of these cell wall inhibitors was not increased (**FIGURE 7a**).

**Figure 7.**
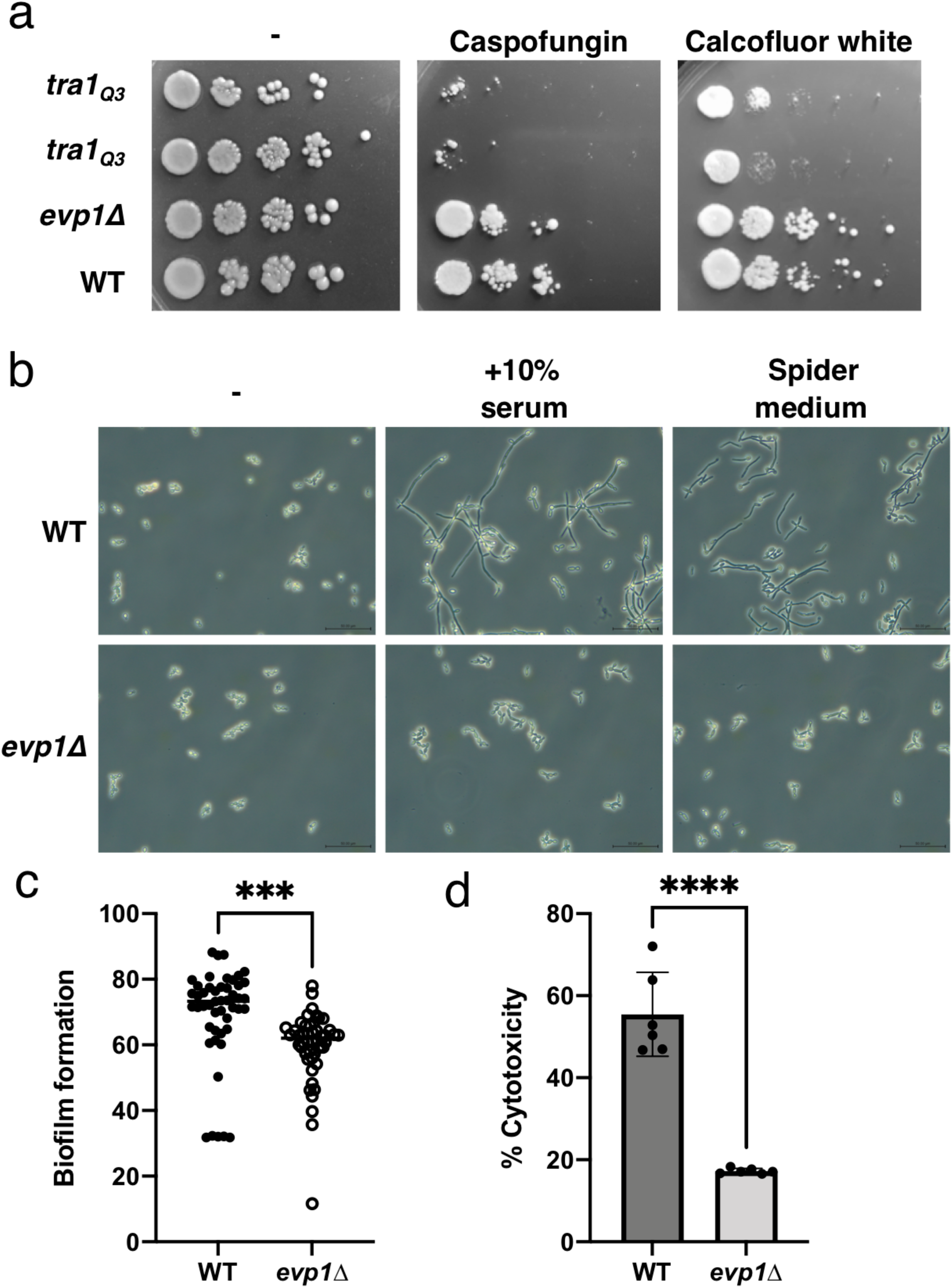
Characterization of the *C. albicans evp1Δ* strain. **(a)** The *C. albicans evp1Δ* strain is not sensitive to cell wall stressors. *C. albicans* strains were serially diluted (1:10 dilutions) and spotted onto solid YPD agar media with or without 0.25 µg/mL caspofungin or 25 µg/mL calcofluor white. **(b)** The *evp1Δ* mutant is impaired in the morphogenetic transition from yeast to filamentous growth in YPD media containing 10% serum or in Spider media. *C. albicans* strains were grown in YPD media alone at 30°C, or in YPD containing 10% serum or in Spider media at 37°C for 4 hours. Cells were observed by bright field microscopy. **(b)** The *evp1Δ* mutant is impaired in biofilm formation relative to a wild-type strain (P < 0.001 (***)). Biofilm growth was quantified by an XTT metabolic readout, measured at OD_490_ and normalized to planktonic growth. Biofilm formation of these fungal strains were measured alongside the *tra1_Q3_* mutants in Figure 6, and thus the wild-type strain measurements are the same between these two figures. **(c)** The *evp1Δ* mutant has reduced macrophage cytotoxicity relative to a wild-type strain (P < 0.0001 (****), error bars represent standard deviation). Macrophage cytotoxicity was quantified by measuring LDH after 6 hours of co-incubation with *C. albicans* strains, as described for Figure 6. Cytotoxicity associated with these fungal strains were measured alongside the *tra1_Q3_* mutants in Figure 6, and thus the wild-strain measurements are the same between these two figures.

Next, we determined whether *EVP1* plays a role in the pathogenesis-associated phenotypes in which *TRA1* has been implicated. Similar to the *tra1_Q3_*, deletion of *EVP1* is severely impaired in the yeast to filamentous growth transition (**FIGURE 7b**), while resulting in no overall growth defect (**FIGURE S1**). In addition, biofilm formation of the *evp1Δ* mutant was reduced relative to wild-type *C. albicans* (**FIGURE 7c,** *P <* 0.0005), and *evp1Δ*-infected macrophages experienced significantly reduced cytotoxicity compared to those infected with wild-type *C. albicans* (**FIGURE 7d,** *P <* 0.0001). Together these results suggest that *EVP1* plays an important role in fungal pathogenesis, though its role in filamentation and biofilm formation is uncoupled from *TRA1*’s role in the resistance to cell wall stressors.

## Discussion

We previously demonstrated that *S. cerevisiae* Tra1_Q3_ is poorly integrated into the SAGA and NuA4 complexes, compromises Tra1-dependent transcription, and mislocalizes to the cytoplasm (47). The transcriptome analysis presented here shows that Tra1 affects multiple cellular processes, in agreement with the role of SAGA as a general cofactor for RNA polymerase II transcription (33, 34, 47, 92). Differential expression of cell wall genes, regulatory association with transcription factors such as Rlm1 and Hsf1, and sensitivity to caspofungin are consistent with the role of Tra1 in stress responses (32), and are in agreement with our previous analysis of other Tra1 derivatives (44, 93).

This work is the first to characterize the role of *TRA1* in the fungal pathogen *C. albicans*. We found that, similar to *S. cerevisiae,* the *C. albicans tra1_Q3_* strain is highly sensitive to cell wall-targeting stressors, including calcofluor white and the antifungal drug caspofungin. Of note, the *tra1_Q3_* allele impairs several *in vitro* measures of fungal pathogenesis, including filamentation, biofilm formation, and macrophage cytotoxicity. Many of these pathogenesis traits are linked, as filamentation is a critical part of biofilm formation and interaction with host immune cells (70, 89, 94). These findings add to a growing literature highlighting the key role of histone acetyltransferases and histone deacetylases in mediating fungal virulence (49–56) and resistance to antifungal drugs (52, 54, 57–61). Recent work with the histone acetyltransferase Gcn5 in *C. albicans* has focused on unravelling the mechanisms by which histone-modifying enzymes alter fungal virulence and sensitivity to antifungal agents, including caspofungin (54). Similar to *TRA1*, loss of *GCN5* in *C. albicans* is associated with defects in filamentation, hypersensitivity to caspofungin, and other virulence and stress-related phenotypes (54). *GCN5*- related caspofungin susceptibility may be linked to Gcn5-mediated regulation of the master transcriptional regulator Efg1 (54), which is involved in caspofungin susceptibility (95). Loss of Gcn5 is associated with altered MAPK signaling, which is involved in remodelling of the fungal cell wall, as well as filamentation and other virulence processes (54, 96–98). Interestingly, Efg1 is one of the transcription factors associated with the genes underpinning the Tra1-regulated caspofungin response (**SUPP FILE 4**). Despite the functional similarities, we find little overlap between our *tra1_Q3_* dataset and the transcriptional profile of *C. albicans* cells depleted of *GCN5* **(FIGURE S3)**. It is difficult to evaluate if these differences emerge from the comparison of a loss-of-function Tra1 mutant with a complete *GCN5* knockout, the role of Tra1 in both SAGA and NuA4, or the possibility that *C. albicans* Gcn5 regulates a subset of genes in the absence of a functional Tra1, as it was recently suggested for *S. cerevisiae* (92). Another interesting result is the absence of *TRA1* upregulation in the *C. albicans tra1_Q3_* strain, which contrasts to what we observed in *S. cerevisiae* (3.2-fold upregulated) (47). Thus, it appears that several genes, including *TRA1,* have undergone a transcriptional rewiring between *S. cerevisiae* and *C. albicans* in response to loss of Tra1 function. Future investigations on the role of *C. albicans* Tra1 in the recruitment of the acetyltrasferases Gcn5 and Esa1 to promoters, as well as Tra1 regulation of histone acetylation, will help clarify these issues. Nevertheless, the key role of these enzymes in fungal virulence and drug susceptibility indicates their potential as antifungal therapeutic targets (99, 100).

The *C. albicans tra1_Q3_* strain had altered phenotypes associated with both antifungal drug resistance, as well as pathogenicity traits such as filamentation. This connection between drug resistance and pathogenesis has been well established amongst pathogenic *Candida* species, and numerous signaling pathways jointly mediate these two important facets of fungal biology (101). One central regulator of fungal pathogenesis and drug resistance is the essential molecular chaperone Hsp90, which plays a key role in *C. albicans* morphogenesis, biofilm formation, resistance to azole and echinocandin antifungals, and virulence in animal models of infection (102–106). Hsp90 itself is regulated by lysine deacetylases including Hos2, Hda1, Rpd3, and Rpd31, which modulate Hsp90’s role in antifungal drug resistance (58) and morphogenesis (107), and numerous downstream cellular pathways are involved in Hsp90- mediated regulation of virulence and drug resistance, including protein kinase A (PKA), and MAPK signaling pathways (102, 108, 109). Recent work in the model yeast *Schizosaccharomyces pombe* found that the Tra1 and Tra2 proteins require Hsp90 along with a cochaperone, the Triple-T complex, for integration into the SAGA and NuA4 complexes (39). If the Tra1-Hsp90 interaction is conserved in *C. albicans*, this interaction may provide at least one rationale for how Hsp90 regulates morphogenesis and virulence.

Transcriptomic analysis revealed *EVP1*, as a previously-uncharacterized gene whose expression is altered in response to caspofungin treatment, and in the *tra1_Q3_* strain. Like *tra1_Q3_*, *evp1Δ* deletion significantly impairs filamentation and biofilm formation; however unlike *tra1_Q3_*, the *evp1Δ* strain retains a wild-type level of resistance to caspofungin, suggesting a more targeted role in fungal pathogenesis. *C. albicans* Evp1 is abundant in extracellular vesicles (EVs) (91). EVs are found in numerous fungal pathogens, including *C. albicans*, *Cryptococcus neoformans*, *Cryptococcus gattii, Histoplasma capsulatum* and *Paracoccidioides brasiliensis*, and play an important role in pathogenesis and host immune cell interactions (110–118). The EVs transport numerous virulence-associated factors to the extracellular environment (113, 116, 117, 119–122). Examples include adhesin proteins and secreted aspartyl protease (SAP) proteins in *C. albicans* EVs (113), and cell wall component glucuronoxylomannan and laccases involved in melanin biosynthesis in EVs from *C. neoformans* (122, 123). Our study demonstrating the involvement of Evp1 in *C. albicans* pathogenesis phenotypes, is consistent with its role as an EV-associated virulence factor.

Our results indicate that Tra1 is an attractive target for combinational antifungal therapy together with current compounds such as caspofungin. Although the exact function of Tra1’s PI3K domain is unknown, it is essential (discussed in more details in Berg *et al.* (47). Like other PIKK family members such as mTor, the activity of the PI3K domain of Tra1 should be druggable. The putative ATP-binding cleft represents an ideal target for small molecule inhibitors and these efforts will be further facilitated by recent Tra1 structural studies (42, 124).

## Materials and Methods

### Yeast maintenance, media, strains

Strains used in this study are listed in Table S1 in the supplemental material, and plasmids are listed in Table S1. Fungal strains (*S. cerevisiae* (S288c derivatives) and *C. albicans*) were cultured on YPD (2% Bacto peptone, 1% yeast extract, 2% glucose) or Synthetic Complete medium (with appropriate amino acids).

### CRISPR design

For CRISPR-based tools used for *C. albicans*, including generating the *tra1_Q3_* mutation, CRISPRi repression and *evp1Δ* deletion, guide RNA N20 sequences were designed based on the efficiency score and specificity using the sgRNA design tool Eukaryotic Pathogen CRISPR gRNA Design Tool (EuPaGDT) (125) available at http://grna.ctegd.uga.edu. *C. albicans* genetic sequences were obtained from the Candida Genome Database (CGD; http://www.candidagenome.org) (90).

### Generation of CRISPR plasmids

CRISPR plasmids for *C. albicans* were generated as previously described (71, 72, 126). CRISPR mutation plasmids targeting *TRA1* or knockout plasmids targeting *EVP1* were engineered in pRS252 (*C. albicans*-optimized Cas9 plasmid (71, 126)). For each construct, two independent sgRNAs were synthesized: for *TRA1*, these sgRNAs flanked the regions containing the three sites being targeted for mutagenesis; for *EVP1*, these sgRNAs flanked the open reading frame (ORF) of the gene. For each plasmid, a gene block (Integrated DNA Technologies, IDT) was synthesized, containing the two sgRNAs, the *SNR52* sgRNA promoter, and regions of homology for Gibson assembly. The *EVP1*-targeting gene block additionally contained regions of homology upstream and downstream of the *EVP1* ORF. These gene blocks were then cloned into the pRS252 backbone using Gibson assembly (127). The pRS252 plasmid was digested with NgoMIV. This digested plasmid was then combined with nuclease-free water, the synthesized gene block fragment (diluted to a concentration of 50 ng/µL), and NEBuilder 2x Hifi DNA Master Mix (New England Biolabs, NEB), for a total volume of 6 μl, and incubated at 50°C for 4 hours. The assembled plasmid was then transformed into chemically competent *Escherichia coli* DH5α using the NEB high efficiency transformation protocol. Plasmid sequences were verified by Sanger sequencing.

CRISPRi repression plasmids targeting *TRA1* for repression were engineered in pRS159 (*C. albicans*-optimized dCas9 plasmid (72)). For this construct, an sgRNA was designed to target dCas9 to the promoter region of *TRA1*. Each sgRNA N20 sequence was obtained from IDT as forward and reverse strands and were reconstituted to 100 µM. Equal volumes of the two complementary oligonucleotides were combined and annealed by heating to 94°C for 2 min and cooling to room temperature. The duplex fragment was cloned into pRS159 using Golden Gate cloning (128). For Golden Gate cloning, the plasmid miniprep, duplexed oligonucleotide, 10x CutSmart buffer, ATP, SapI, T4 DNA ligase, and nuclease-free water were combined and the mixture was incubated in a thermocycler under the following cycling conditions: (37°C, 2 min; 16°C, 5 min) for 99 cycles; 65°C, 15 min; 80°C, 15 min. After cycling was complete, 1 µl of SapI enzyme was added to each reaction, and incubated at 37°C for 1 hour. Buffers, enzymes, and ATP were obtained from NEB. Plasmids were verified by Sanger sequencing.

### C. albicans transformation

*C. albicans* cells were transformed as previously described (71). Briefly, the plasmids to be transformed were miniprepped and plasmid quality and purity were determined using a Nanodrop spectrophotometer (Tecan). Cas9 and dCas9 plasmids were linearized with the PacI restriction enzyme (NEB) to enable plasmid integration at the *NEUT5L* locus. For CRISPRi, linearized plasmid alone was transformed into *C. albicans*. Other CRISPR plasmids were co-transformed with repair templates (gRS11 for *tra1*_Q3_ containing Q3 mutations, gRS76 for *evp1Δ*). Linearized plasmids, repair templates, and *C. albicans* cells were incubated with 50% polyethylene glycol, 10x Tris-EDTA buffer, 1 M lithium acetate (pH 7.4), salmon sperm DNA, and 2 M dithiothreitol and incubated at 30°C for 1 hour and at 42°C for 45 min. Post-transformation, cells were grown in YPD medium for 4 hours at 30°C with shaking and transformants were selected for on YPD agar plates containing 200 µg/ml nourseothricin (Jena Biosciences). *C. albicans* transformed colonies were PCR-verified for presence of the Cas9 or dCas9 construct and/or specific gene deletions. The presence of the three targeted point mutations (R3471Q, R3472Q, and R3538Q) in *TRA1* was confirmed through Sanger sequencing.

### CFU spot platings

Wild-type and *tra1_Q3_* strains were grown in 5 mL YPD overnight at 30°C. Strains were then diluted to an OD_600_ of 0.05 and grown to early log phase, with an OD_600_ of 0.2, by shaking at 30°C for 2-3 hours. 100μl of each strain was transferred to the first well of each row of a 96-well plate and 10-fold serial dilutions made in YPD. 5 μl of each row was spotted onto plain YPD plates, or YPD plates containing 25 μg/ml calcofluor white or 0.25μg/ml caspofungin. Each plate was set up in triplicate. Plates were incubated at 30°C for 2 days.

### RNA preparation and sequencing

For *C. albicans*, RNA samples were generated in triplicate for each mutant. The following triplicate sets were grown overnight in 50 mL inoculated YPD tubes, for a total of 12 tubes: wild type without drug, wild type grown with 10 ng/ml caspofungin, *tra1_Q3_* mutant without drug, and *tra1_Q3_* mutant grown with 10ng/ml caspofungin. The appropriate samples were inoculated with drug when at an OD_600_ of 0.1, and then grown to an OD_600_ of 0.2 with drug for approximately 2 hours at 30°C. RNA samples were then extracted using the Presto Mini RNA Yeast Kit (Geneaid). For *S. cerevisiae*, RNA was extracted from three replicates from each strain at early log phase growth in synthetic complete medium. RNA was extracted using the MasterPure Yeast RNA purification kit (Lucigen) according to the manufacturer’s instructions. Sample purity was checked using a tapestation bioanalyzer to ensure a RIN value of 8.0 or higher before being vacuum dried. Dried sample pellets were sent to Genewiz for further analysis.

Total RNA sequencing was performed by Genewiz (South Plainfield, NJ). Stranded Illumina TruSeq cDNA libraries with poly dT enrichment were prepared from high quality total RNA (RIN > 8). Libraries were sequenced on an Illumina HiSeq, yielding between 26.4 - 37.9 million 150 bp paired end sequencing reads per sample. The data have been deposited in NCBI’s Gene Expression Omnibus (129) and are accessible through GEO Series accession number GSE168955 for *S. cerevisiae* (https://www.ncbi.nlm.nih.gov/geo/query/acc.cgi?acc=GSE168955) and GSE168988 for *C. albicans* (https://www.ncbi.nlm.nih.gov/geo/query/acc.cgi?acc=GSE168988).

### Quality control, trimming, read alignment and differential gene expression analysis

FASTQ files were analyzed using a customized bioinformatics workflow. Adapter sequences and low quality bases were trimmed using the default settings of Trimmomatic (130). Sequence quality was analyzed using FastQC (http://www.bioinformatics.babraham.ac.uk/projects/fastqc/). Sequence reads were aligned to the *Saccharomyces cerevisiae S288C* reference genome (assembly R64-2-1; https://www.yeastgenome.org/) or the *Candida albicans* SC5314 reference genome (assembly 21; http://www.candidagenome.org) using STAR (131). Only reads that uniquely mapped to the reference genome were kept. Read counts for each gene were summarized using featureCounts (132). Differential expression analysis was performed using the DESeq2 R package (133) using a custom R script (**SUPP FILE 5**) with a Benjamini-Hochberg adjusted *P*-value cut-off of <= 0.05. The volcano plot presented in **Figure S3** was generated using VolcaNoseR (134).

### Filamentation assay and microscopy

Wild-type and mutant strains were grown overnight in 5 mL of YPD at 30°C. OD_600_ reads were taken and strain growth normalized by diluting samples to OD_600_ = 0.2. Cells were subcultured and incubated in YPD with 10% serum or in Spider medium to induce filamentation. Strains induced with serum or grown in Spider medium were incubated at 37°C, while strains grown in non-filamentous conditions were incubated at 30°C, for four hours. Filamentation was observed via bright field microscopy.

### Growth curves

Overnight cultures of 5 mL YPD inoculated with *C. albicans* were grown at 30°C, with shaking at 250 rpm. Growth of the strains was evaluated by measuring optical density at 600 nm and strain growth was normalized by diluting to OD_600_ = 0.1. In a 96-well plate containing 100 μL of fresh YPD, 100 μL of each sample was added, followed by the addition of 50 μL of mineral oil to prevent evaporation. OD_600_ was determined every 15 min for 24 hours in a microplate reader, with shaking between reads. The data was graphed using Graphpad Prism version 8.

### Biofilm growth

Wild-type, *tra1_Q3_*, and *evp1Δ* strains were incubated overnight in 5 mL YPD at 37°C and 250 rpm. The cultures were normalized to the same OD_600_ by diluting with RPMI liquid medium. 100 μL of each subcultured strain was distributed into flat bottom 96-well polystyrene plates containing 100 μL of RPMI in each well. A row containing 200 μL of RPMI was used as a negative control. The 96-well plates were wrapped in aluminum foil, then incubated at 37°C for 72 hours. 120 μL of planktonic cells was removed from each well, dispensed into new 96-well plates, and OD_600_ determined in a microplate reader. The biofilm plates were washed twice with 200 μL of 1x PBS buffer and dried in a fume hood for ∼1 hour. Next, 90 μL of XTT (1 mg/mL) and 10 μL of PMS (0.32 mg/mL) were added sequentially to every well, the plates were wrapped in aluminum foil, and incubated for 2 hours at 30°C. OD_490_ was determined in a microplate reader and the XTT biofilm growth values normalized to planktonic cell growth.

### Macrophage cell lines

Immortalized macrophages, of BALB/c murine origin, were used to conduct macrophage infection assays. Prior to infection, cells were passaged into three 150 mm tissue culture dishes of fresh DMEM (Gibco) medium, supplemented with 2 mM L-glutamine (Gibco), 2 mM penicillin-streptomycin (Lonza), and 10% heat-inactivated fetal bovine serum (Life Technologies). These cells were then incubated at 37°C and 5% CO_2_, for three days to grow up to 70-80% confluence. On infection day, cells were washed and collected in DMEM medium lacking 2 mM penicillin-streptomycin (- pen - strep). The concentration of living cells was determined using the Countess™ II automated cell counter (Invitrogen). The appropriate volume of cells were added to each well of a 12-well tissue culture plate, to obtain 500,000 cells in 1 mL medium. Cells were incubated for 3 hours at 37°C and 5% CO_2_, to allow adherence to the plate surface prior to infection.

### C. albicans-macrophage infection

Overnight cultures of *C. albicans* strains were grown in 5 mL YPD at 37°C and 250 rpm, the day before infection. Cells were then subcultured in 5 mL of fresh YPD and grown to an OD_600_ of 1.0. Cells were washed and resuspended in 1 mL DMEM (- pen - strep). Cell count was determined on a hemocytometer under a light microscope. *C. albicans* strains were added to macrophages at a ratio of 1 yeast cell: 10 macrophages. Assuming 500,000 macrophages are in each well of the tissue culture plate, the appropriate volume of *C. albicans* cells was added to obtain 50,000 cells/well. DMEM (- pen - strep) medium, as well as uninfected macrophages, were included as controls.

*C. albicans* and macrophages were co-incubated for 1 hour at 37°C and 5% CO_2_ to allow infection. All further incubations were at 37°C and 5% CO_2_. Wells were washed with 1x sterile PBS (without calcium or magnesium, Lonza) and cells incubated 30 min with DMEM (- pen - strep), supplemented with 5 µg/mL caspofungin to kill residual *C. albicans*. Wells were washed three times with 1x PBS and provided with 1 mL of DMEM (- pen - strep) medium. Plates were incubated and the cytotoxicity of each *C. albicans* strain were measured using the LDH assay at the 1-hour, 3-hour, 6-hour, and 18-hour time points. The LDH assay was conducted using the CytoTox 96® Non-Radioactive Cytotoxicity Assay kit (Promega). At each time point, the supernatant from each well was collected and 1 mL of 1.2% Triton X (diluted with PBS) was added to lyse cells for 30 minutes in the dark. On a 96-well plate, 50 uL of LDH substrate was added to 50 µL of both the supernatant and cell lysate for each treatment condition, as well as the untreated and media control conditions. Plates were incubated in the dark for 30 minutes, after which 50 µL of the stop solution was added. The OD_490_ was measured for each condition, and the % cytotoxicity of each strain (treatment) was calculated using these OD_490_ values in the following equation: % cytotoxicity = (treatment - media control)/(uninfected control - media control) x 100%.

### Data availability

The data from the RNA-seq experiments have been deposited in NCBI’s Gene Expression Omnibus (129) and are accessible through GEO Series accession number GSE168955 for *S. cerevisiae* (https://www.ncbi.nlm.nih.gov/geo/query/acc.cgi?acc=GSE168955) and GSE168988 for *C. albicans* (https://www.ncbi.nlm.nih.gov/geo/query/acc.cgi?acc=GSE168988). *C. albicans* CRISPR plasmid backbones are available via Addgene: Cas9 plasmid Addgene ID: 89576; CRISPRi dCas9 plasmid Addgene ID: 122378.

## Acknowledgments

This work was supported by a CIHR Project Grant (PJT 162195) and NSERC Discovery Grant (RGPIN-2018-4914) to RSS. PL is supported by a NSERC Discovery Grant (RGPIN-2015-06400) and a Canadian Foundation for Innovation (CFI) John R. Evans Leader Fund Grant (65183). CJB is supported by a NSERC Discovery Grant (RGPIN-2015-04394). YJ was supported by a Masters to PhD Transfer Scholarship from the Dean of the Schulich Faculty of Medicine and Dentistry at Western University. MDB was supported by an NSERC Alexander Graham Bell Canada Graduate Scholarship.

## Figure legends

**Figure S1. *EVP1* and *TRA1* compromised strains are not defective in overall growth rate**. Kinetic growth curves between *C. albicans* wild-type strain, *evp1Δ* mutant and *tra1_Q3_* mutant strains indicate no differences in overall growth dynamics over time.

**Figure S2. *EVP1* is differentially regulated in *tra1_Q3_* mutant strains in the presence or absence of caspofungin**. Normalized counts for *EVP1* from RNA-seq analysis between the *C. albicans* wild type and *tra1_Q3_* mutant strains in the presence or absence of caspofungin (not significant (n.s.), P < 0.05 (*); Welch’s *t*-test).

**Figure S3. Limited overlap between the Tra1- and Gcn5-regulated genes in *C. albicans*. (a)** Volcano plot showing the distribution of the previously identified 336 differentially expressed genes in *gcn5Δ (*from Shivarathri *et al.)* within the *tra1_Q3_* dataset. Genes significantly (log_2_ Fold Change *tra1_Q3_*/*TRA1* > 1 and *P* < 0.05) and highlighted in blue (downregulated) and red (upregulated). **(b)** Differentially regulated Gcn5 targets in *tra1_Q3_* dataset are listed in the table with both the log_2_ fold change within the original *gcn5Δ* and the *tra1_Q3_* datasets.

## References

1. Bongomin F, Gago S, Oladele R, Denning D. 2017. Global and multi-national prevalence of fungal diseases—estimate precision. Journal of Fungi 3:57.

2. Brown GD, Denning DW, Gow NAR, Levitz SM, Netea MG, White TC. 2012. Hidden killers: human fungal infections. Sci Transl Med 4:165rv13.

3. Pfaller MA, Diekema DJ. 2007. Epidemiology of invasive candidiasis: a persistent public health problem. Clin Microbiol Rev 20:133–163.

4. Kullberg BJ, Arendrup MC. 2015. Invasive candidiasis. N Engl J Med 373:1445–1456.

5. Perfect JR. 2017. The antifungal pipeline: a reality check. Nat Rev Drug Discov 16:603– 616.

6. Lee Y, Puumala E, Robbins N, Cowen LE. 2020. Antifungal drug resistance: molecular mechanisms in *Candida albicans* and beyond. Chem Rev https://doi.org/10.1021/acs.chemrev.0c00199.

7. Cowen LE, Anderson JB, Kohn LM. 2002. Evolution of drug resistance in *Candida albicans*. Annu Rev Microbiol 56:139–165.

8. Geddes-McAlister J, Shapiro RS. 2019. New pathogens, new tricks: emerging, drug-resistant fungal pathogens and future prospects for antifungal therapeutics. Ann N Y Acad Sci 1435:57–78.

9. Miceli MH, Díaz JA, Lee SA. 2011. Emerging opportunistic yeast infections. Lancet Infect Dis 11:142–151.

10. Whaley SG, Berkow EL, Rybak JM, Nishimoto AT, Barker KS, Rogers PD. 2016. Azole antifungal resistance in *Candida albicans* and emerging non-*albicans Candida* species. Front Microbiol 7:2173.

11. Chowdhary A, Sharma C, Meis JF. 2017. *Candida auris*: A rapidly emerging cause of hospital-acquired multidrug-resistant fungal infections globally. PLoS Pathog 13:e1006290.

12. Aguilar-Zapata D, Petraitiene R, Petraitis V. 2015. Echinocandins: The expanding antifungal armamentarium. Clin Infect Dis 6:S604–11.

13. Kurtz MB, Douglas CM. 1997. Lipopeptide inhibitors of fungal glucan synthase. Medical Mycology 35:79–86

14. Pound MW, Townsend ML, Drew RH. 2010. Echinocandin pharmacodynamics: review and clinical implications. J Antimicrob Chemother 65:1108–1118.

15. Cleveland AA, Farley MM, Harrison LH, Stein B, Hollick R, Lockhart SR, Magill SS, Derado G, Park BJ, Chiller TM. 2012. Changes in incidence and antifungal drug resistance in candidemia: results from population-based laboratory surveillance in Atlanta and Baltimore, 2008-2011. Clin Infect Dis 55:1352–1361.

16. Perlin DS. 2015. Echinocandin resistance in *Candida*. Clin Infect Dis 61 Suppl 6:S612–7.

17. Pappas PG, Kauffman CA, Andes DR, Clancy CJ, Marr KA, Ostrosky-Zeichner L, Reboli AC, Schuster MG, Vazquez JA, Walsh TJ, Zaoutis TE, Sobel JD. 2016. Clinical practice guideline for the management of candidiasis: 2016 Update by the Infectious Diseases Society of America. Clin Infect Dis 62:e1–50.

18. Alexander BD, Johnson MD, Pfeiffer CD, Jiménez-Ortigosa C, Catania J, Booker R, Castanheira M, Messer SA, Perlin DS, Pfaller MA. 2013. Increasing echinocandin resistance in *Candida glabrata*: clinical failure correlates with presence of *FKS* mutations and elevated minimum inhibitory concentrations. Clin Infect Dis 56:1724–1732.

19. Garcia-Effron G, Chua DJ, Tomada JR, DiPersio J, Perlin DS, Ghannoum M, Bonilla H. 2010. Novel *FKS* mutations associated with echinocandin resistance in *Candida* species. Antimicrob Agents Chemother 54:2225–2227.

20. Cowen LE, Steinbach WJ. 2008. Stress, drugs, and evolution: the role of cellular signaling in fungal drug resistance. Eukaryot Cell 7:747–764.

21. Víglaš J, Olejníková P. 2020. Signalling mechanisms involved in stress response to antifungal drugs. Res Microbiol https://doi.org/10.1016/j.resmic.2020.10.001.

22. Cannon RD, Lamping E, Holmes AR, Niimi K, Tanabe K, Niimi M, Monk BC. 2007. *Candida albicans* drug resistance another way to cope with stress. Microbiology 153:3211–3217.

23. Shor E, Perlin DS. 2015. Coping with stress and the emergence of multidrug resistance in fungi. PLoS Pathog 11:e1004668.

24. Yun M, Wu J, Workman JL, Li B. 2011. Readers of histone modifications. Cell Res 21:564– 578.

25. Bannister AJ, Kouzarides T. 2011. Regulation of chromatin by histone modifications. Cell Res 21:381–395.

26. Grant PA, Duggan L, Côté J, Roberts SM, Brownell JE, Candau R, Ohba R, Owen-Hughes T, Allis CD, Winston F, Berger SL, Workman JL. 1997. Yeast Gcn5 functions in two multisubunit complexes to acetylate nucleosomal histones: characterization of an Ada complex and the SAGA (Spt/Ada) complex. Genes Dev 11:1640–1650.

27. Allard S, Utley RT, Savard J, Clarke A, Grant P, Brandl CJ, Pillus L, Workman JL, Côté J. 1999. NuA4, an essential transcription adaptor/histone H4 acetyltransferase complex containing Esa1p and the ATM-related cofactor Tra1p. EMBO J 18:5108–5119.

28. Auger A, Galarneau L, Altaf M, Nourani A, Doyon Y, Utley RT, Cronier D, Allard S, Côté J. 2008. Eaf1 is the platform for NuA4 molecular assembly that evolutionarily links chromatin acetylation to ATP-dependent exchange of histone H2A variants. Mol Cell Biol 28:2257– 2270.

29. Choudhary C, Kumar C, Gnad F, Nielsen ML, Rehman M, Walther TC, Olsen JV, Mann M. 2009. Lysine acetylation targets protein complexes and co-regulates major cellular functions. Science 325:834–840.

30. Henriksen P, Wagner SA, Weinert BT, Sharma S, Bacinskaja G, Rehman M, Juffer AH, Walther TC, Lisby M, Choudhary C. 2012. Proteome-wide analysis of lysine acetylation suggests its broad regulatory scope in *Saccharomyces cerevisiae*. Mol Cell Proteomics 11:1510–1522.

31. Mitchell L, Huard S, Cotrut M, Pourhanifeh-Lemeri R, Steunou A-L, Hamza A, Lambert J-P, Zhou H, Ning Z, Basu A, Côté J, Figeys DA, Baetz K. 2013. mChIP-KAT-MS, a method to map protein interactions and acetylation sites for lysine acetyltransferases. Proc Natl Acad Sci U S A 110:E1641–50.

32. Huisinga KL, Pugh BF. 2004. A genome-wide housekeeping role for TFIID and a highly regulated stress-related role for SAGA in *Saccharomyces cerevisiae*. Mol Cell 13:573–585.

33. Bonnet J, Wang C-Y, Baptista T, Vincent SD, Hsiao W-C, Stierle M, Kao C-F, Tora L, Devys D. 2014. The SAGA coactivator complex acts on the whole transcribed genome and is required for RNA polymerase II transcription. Genes Dev 28:1999–2012.

34. Baptista T, Grünberg S, Minoungou N, Koster MJE, Marc Timmers HT, Hahn S, Devys D, Tora L. 2018. SAGA is a general cofactor for RNA polymerase II transcription. Molecular Cell 68:130–143

35. Grant PA, Schieltz D, Pray-Grant MG, Yates JR 3rd, Workman JL. 1998. The ATM-related cofactor Tra1 is a component of the purified SAGA complex. Mol Cell 2:863–867.

36. Saleh A, Schieltz D, Ting N, McMahon SB, Litchfield DW, Yates JR 3rd, Lees-Miller SP, Cole MD, Brandl CJ. 1998. Tra1p is a component of the yeast Ada.Spt transcriptional regulatory complexes. J Biol Chem 273:26559–26565.

37. Elías-Villalobos A, Fort P, Helmlinger D. 2019. New insights into the evolutionary conservation of the sole PIKK pseudokinase Tra1/TRRAP. Biochem Soc Trans 47:1597– 1608.

38. Lempiäinen H, Halazonetis TD. 2009. Emerging common themes in regulation of PIKKs and PI3Ks. EMBO J 28:3067–3073.

39. Elías-Villalobos A, Toullec D, Faux C, Séveno M, Helmlinger D. 2019. Chaperone-mediated ordered assembly of the SAGA and NuA4 transcription co-activator complexes in yeast. Nat Comm 10:5237

40. Genereaux J, Kvas S, Dobransky D, Karagiannis J, Gloor GB, Brandl CJ. 2012. Genetic evidence links the ASTRA protein chaperone component Tti2 to the SAGA transcription factor Tra1. Genetics 191:765–780.

41. Baretić D, Pollard HK, Fisher DI, Johnson CM, Santhanam B, Truman CM, Kouba T, Fersht AR, Phillips C, Williams RL. 2017. Structures of closed and open conformations of dimeric human ATM. Sci Adv 3:e1700933.

42. Díaz-Santín LM, Lukoyanova N, Aciyan E, Cheung AC. 2017. Cryo-EM structure of the SAGA and NuA4 coactivator subunit Tra1 at 3.7 angstrom resolution. eLife 6:e28384

43. Knutson BA, Hahn S. 2011. Domains of Tra1 important for activator recruitment and transcription coactivator functions of SAGA and NuA4 complexes. Mol Cell Biol 31:818– 831.

44. Mutiu AI, Hoke SMT, Genereaux J, Hannam C, MacKenzie K, Jobin-Robitaille O, Guzzo J, Côté J, Andrews B, Haniford DB, Brandl CJ. 2007. Structure/function analysis of the phosphatidylinositol-3-kinase domain of yeast tra1. Genetics 177:151–166.

45. Lovejoy CA, Cortez D. 2009. Common mechanisms of PIKK regulation. DNA Repair 8:1004–1008.

46. McMahon SB, Van Buskirk HA, Dugan KA, Copeland TD, Cole MD. 1998. The novel ATM-related protein TRRAP is an essential cofactor for the c-Myc and E2F oncoproteins. Cell 94:363–374.

47. Berg MD, Genereaux J, Karagiannis J, Brandl CJ. 2018. The pseudokinase domain of *Saccharomyces cerevisiae* Tra1 is required for nuclear localization and incorporation into the SAGA and NuA4 Complexes. G3 8:1943–1957

48. O’Kane CJ, Weild R, M Hyland E. 2020. Chromatin Structure and Drug Resistance in *Candida* spp. J Fungi 6:121

49. Hnisz D, Majer O, Frohner IE, Komnenovic V, Kuchler K. 2010. The Set3/Hos2 histone deacetylase complex attenuates cAMP/PKA signaling to regulate morphogenesis and virulence of *Candida albicans*. PLoS Pathog 6:e1000889.

50. Hnisz D, Bardet AF, Nobile CJ, Petryshyn A, Glaser W, Schöck U, Stark A, Kuchler K. 2012. A histone deacetylase adjusts transcription kinetics at coding sequences during *Candida albicans* morphogenesis. PLoS Genet 8:e1003118.

51. Zacchi LF, Schulz WL, Davis DA. 2010. *HOS2* and *HDA1* encode histone deacetylases with opposing roles in *Candida albicans* morphogenesis. PLoS One 5:e12171.

52. Nobile CJ, Fox EP, Hartooni N, Mitchell KF, Hnisz D, Andes DR, Kuchler K, Johnson AD. 2014. A histone deacetylase complex mediates biofilm dispersal and drug resistance in *Candida albicans*. mBio 5:e01201–14.

53. Wang X, Chang P, Ding J, Chen J. 2013. Distinct and redundant roles of the two MYST histone acetyltransferases Esa1 and Sas2 in cell growth and morphogenesis of *Candida albicans*. Eukaryot Cell 12:438–449.

54. Shivarathri R, Tscherner M, Zwolanek F, Singh NK, Chauhan N, Kuchler K. 2019. The fungal histone acetyl transferase Gcn5 controls virulence of the human pathogen *Candida albicans* through Multiple Pathways. Sci Rep 9:9445.

55. Tscherner M, Zwolanek F, Jenull S, Sedlazeck FJ, Petryshyn A, Frohner IE, Mavrianos J, Chauhan N, von Haeseler A, Kuchler K. 2015. The *Candida albicans* histone acetyltransferase Hat1 regulates stress resistance and virulence via distinct chromatin assembly pathways. PLoS Pathog 11:e1005218.

56. da Rosa JL, Boyartchuk VL, Zhu LJ, Kaufman PD. 2010. Histone acetyltransferase Rtt109 is required for *Candida albicans* pathogenesis. Proc Natl Acad Sci U S A 107:1594–1599.

57. Smith WL, Edlind TD. 2002. Histone deacetylase inhibitors enhance *Candida albicans* sensitivity to azoles and related antifungals: correlation with reduction in *CDR* and *ERG* upregulation. Antimicrob Agents Chemother 46:3532–3539.

58. Robbins N, Leach MD, Cowen LE. 2012. Lysine deacetylases Hda1 and Rpd3 regulate Hsp90 function thereby governing fungal drug resistance. Cell Rep 2:878–888.

59. Ramírez-Zavala B, Mogavero S, Schöller E, Sasse C, Rogers PD, Morschhäuser J. 2014. SAGA/ADA complex subunit Ada2 is required for Cap1- but not Mrr1-mediated upregulation of the *Candida albicans* multidrug efflux pump *MDR1*. Antimicrob Agents Chemother 58:5102–5110.

60. Sellam A, Askew C, Epp E, Lavoie H, Whiteway M, Nantel A. 2009. Genome-wide mapping of the coactivator Ada2p yields insight into the functional roles of SAGA/ADA complex in *Candida albicans*. Mol Biol Cell 20:2389–2400.

61. Yu S-J, Chang Y-L, Chen Y-L. 2018. Deletion of *ADA2* increases antifungal drug susceptibility and virulence in *Candida glabrata*. Antimicrob Agents Chemother 62.

62. Mitchell L, Lambert J-P, Gerdes M, Al-Madhoun AS, Skerjanc IS, Figeys D, Baetz K. 2008. Functional dissection of the NuA4 histone acetyltransferase reveals its role as a genetic hub and that Eaf1 is essential for complex integrity. Mol Cell Biol 28:2244–2256.

63. Helmlinger D, Marguerat S, Villén J, Swaney DL, Gygi SP, Bähler J, Winston F. 2011. Tra1 has specific regulatory roles, rather than global functions, within the SAGA co-activator complex. EMBO J 30:2843–2852.

64. Levin DE. 2011. Regulation of cell wall biogenesis in *Saccharomyces cerevisiae*: the cell wall integrity signaling pathway. Genetics 189:1145–1175.

65. Sanz AB, García R, Rodríguez-Peña JM, Nombela C, Arroyo J. 2016. Cooperation between SAGA and SWI/SNF complexes is required for efficient transcriptional responses regulated by the yeast MAPK Slt2. Nucleic Acids Res 44:7159–7172.

66. Hoke SMT, Irina Mutiu A, Genereaux J, Kvas S, Buck M, Yu M, Gloor GB, Brandl CJ. 2010. Mutational analysis of the C-terminal FATC domain of *Saccharomyces cerevisiae* Tra1. Curr Genet 56:447–465.

67. Monteiro PT, Oliveira J, Pais P, Antunes M, Palma M, Cavalheiro M, Galocha M, Godinho CP, Martins LC, Bourbon N, Mota MN, Ribeiro RA, Viana R, Sá-Correia I, Teixeira MC. 2020. YEASTRACT+: a portal for cross-species comparative genomics of transcription regulation in yeasts. Nucleic Acids Res 48:D642–D649.

68. Lempiäinen H, Uotila A, Urban J, Dohnal I, Ammerer G, Loewith R, Shore D. 2009. Sfp1 interaction with TORC1 and Mrs6 reveals feedback regulation on TOR signaling. Mol Cell 33:704–716.

69. Jiang Y, Berg MD, Genereaux J, Ahmed K, Duennwald ML, Brandl CJ, Lajoie P. 2019. Sfp1 links TORC1 and cell growth regulation to the yeast SAGA-complex component Tra1 in response to polyQ proteotoxicity. Traffic 20:267–283.

70. Shapiro RS, Robbins N, Cowen LE. 2011. Regulatory circuitry governing fungal development, drug resistance, and disease. Microbiol Mol Biol Rev 75:213–267.

71. Halder V, Porter CBM, Chavez A, Shapiro RS. 2019. Design, execution, and analysis of CRISPR–Cas9-based deletions and genetic interaction networks in the fungal pathogen *Candida albicans*. Nat Protoc 14:955–975.

72. Wensing L, Sharma J, Uthayakumar D, Proteau Y, Chavez A, Shapiro RS. 2019. A CRISPR interference platform for efficient genetic repression in *Candida albicans*. mSphere 4.

73. Barbaric S, Reinke H, Hörz W. 2003. Multiple mechanistically distinct functions of SAGA at the *PHO5* promoter. Mol Cell Biol 23:3468–3476.

74. Adkins MW, Williams SK, Linger J, Tyler JK. 2007. Chromatin disassembly from the *PHO5* promoter is essential for the recruitment of the general transcription machinery and coactivators. Mol Cell Biol 27:6372–6382.

75. Nourani A, Utley RT, Allard S, Côté J. 2004. Recruitment of the NuA4 complex poises the PHO5 promoter for chromatin remodeling and activation. EMBO J 23:2597–2607.

76. Murad AMA. 2001. *NRG1* represses yeast-hypha morphogenesis and hypha-specific gene expression in *Candida albicans*. EMBO J 20:4742–52

77. Cleary IA, Saville SP. 2010. An analysis of the impact of *NRG1* overexpression on the *Candida albicans* response to specific environmental stimuli. Mycopathologia 170:1–10

78. Moran GP, MacCallum DM, Spiering MJ, Coleman DC, Sullivan DJ. 2007. Differential regulation of the transcriptional repressor *NRG1* accounts for altered host-cell interactions in *Candida albicans* and *Candida dubliniensis*. Mol Microbiol 66:915–29

79. Urrialde V, Prieto D, Pla J, Alonso-Monge R. 2016. The *Candida albicans* Pho4 transcription factor mediates susceptibility to stress and influences fitness in a mouse commensalism model. Front Microbiol 7:1062.

80. Song Y-D, Hsu C-C, Lew SQ, Lin C-H. 2020. *Candida tropicalis RON1* is required for hyphal formation, biofilm development, and virulence but is dispensable for N-acetylglucosamine catabolism. Med Mycol https://doi.org/10.1093/mmy/myaa063.

81. Chandy M, Gutiérrez JL, Prochasson P, Workman JL. 2006. SWI/SNF displaces SAGA-acetylated nucleosomes. Eukaryot Cell 5:1738–1747.

82. Dewhurst-Maridor G, Abegg D, David FPA, Rougemont J, Scott CC, Adibekian A, Riezman H. 2017. The SAGA complex, together with transcription factors and the endocytic protein Rvs167p, coordinates the reprofiling of gene expression in response to changes in sterol composition in. Mol Biol Cell 28:2637–2649.

83. Lin L, Chamberlain L, Zhu LJ, Green MR. 2012. Analysis of Gal4-directed transcription activation using Tra1 mutants selectively defective for interaction with Gal4. Proc Natl Acad Sci U S A 109:1997–2002.

84. Proft M, Struhl K. 2002. Hog1 kinase converts the Sko1-Cyc8-Tup1 repressor complex into an activator that recruits SAGA and SWI/SNF in response to osmotic stress. Mol Cell 9:1307–17

85. Bruno VM, Kalachikov S, Subaran R, Nobile CJ, Kyratsous C, Mitchell AP. 2006. Control of the *C. albicans* cell wall damage response by transcriptional regulator Cas5. PLoS Pathog 2:e21.

86. Heredia MY, Ikeh MAC, Gunasekaran D, Conrad KA, Filimonava S, Marotta DH, Nobile CJ, Rauceo JM. 2020. An expanded cell wall damage signaling network is comprised of the transcription factors Rlm1 and Sko1 in *Candida albicans*. PLoS Genet 16:e1008908.

87. Heredia MY, Gunasekaran D, Ikeh MAC, Nobile CJ, Rauceo JM. 2020. Transcriptional regulation of the caspofungin-induced cell wall damage response in *Candida albicans*. Curr Genet 66:1059–1068.

88. Lohse MB, Gulati M, Johnson AD, Nobile CJ. 2018. Development and regulation of single- and multi-species *Candida albicans* biofilms. Nat Rev Microbiol 16:19–31.

89. Gow NAR, van de Veerdonk FL, Brown AJP, Netea MG. 2011. *Candida albicans* morphogenesis and host defence: discriminating invasion from colonization. Nat Rev Microbiol 10:112–122.

90. Skrzypek MS, Binkley J, Binkley G, Miyasato SR, Simison M, Sherlock G. 2017. The *Candida* Genome Database (CGD): incorporation of Assembly 22, systematic identifiers and visualization of high throughput sequencing data. Nucleic Acids Res 45:D592–D596.

91. Dawson CS, Garcia-Ceron D, Rajapaksha H, Faou P, Bleackley MR, Anderson MA. 2020. Protein markers for *Candida albicans* EVs include claudin-like Sur7 family proteins. J Extracell Vesicles 9:1750810.

92. Bruzzone MJ, Grünberg S, Kubik S, Zentner GE, Shore D. 2018. Distinct patterns of histone acetyltransferase and Mediator deployment at yeast protein-coding genes. Genes Dev 32:1252–1265.

93. Hoke SMT, Guzzo J, Andrews B, Brandl CJ. 2008. Systematic genetic array analysis links the *Saccharomyces cerevisiae* SAGA/SLIK and NuA4 component Tra1 to multiple cellular processes. BMC Genet 9:46.

94. Whiteway M, Bachewich C. 2007. Morphogenesis in *Candida albicans*. Annu Rev Microbiol 61:529–553.

95. Gregori C, Glaser W, Frohner IE, Reinoso-Martín C, Rupp S, Schüller C, Kuchler K. 2011. Efg1 Controls caspofungin-induced cell aggregation of *Candida albicans* through the adhesin Als1. Eukaryot Cell 10:1694–1704.

96. Monge RA, Román E, Nombela C, Pla J. 2006. The MAP kinase signal transduction network in *Candida albicans*. Microbiology 152:905–912.

97. Román E, Arana DM, Nombela C, Alonso-Monge R, Pla J. 2007. MAP kinase pathways as regulators of fungal virulence. Trends Microbiol 15:181–190.

98. Chow J, Notaro M, Prabhakar A, Free SJ, Cullen PJ. 2018. Impact of fungal MAPK pathway targets on the cell wall. J Fungi 4:93

99. Bauer I, Varadarajan D, Pidroni A, Gross S, Vergeiner S, Faber B, Hermann M, Tribus M, Brosch G, Graessle S. 2016. A Class 1 histone deacetylase with potential as an antifungal target. mBio 7: e00831–16

100. Kuchler K, Jenull S, Shivarathri R, Chauhan N. 2016. Fungal KATs/KDACs: a new highway to better antifungal drugs? PLoS Pathog 12:e1005938.

101. Sharma J, Rosiana S, Razzaq I, Shapiro RS. 2019. Linking cellular morphogenesis with antifungal treatment and susceptibility in *Candida* pathogens. J Fungi 5:17

102. Shapiro RS, Uppuluri P, Zaas AK, Collins C, Senn H, Perfect JR, Heitman J, Cowen LE. 2009. Hsp90 orchestrates temperature-dependent *Candida albicans* morphogenesis via Ras1-PKA signaling. Curr Biol 19:621–629.

103. Cowen LE, Lindquist S. 2005. Hsp90 potentiates the rapid evolution of new traits: drug resistance in diverse fungi. Science 309:2185–2189.

104. Singh SD, Robbins N, Zaas AK, Schell WA, Perfect JR, Cowen LE. 2009. Hsp90 governs echinocandin resistance in the pathogenic yeast *Candida albicans* via calcineurin. PLoS Pathog 5:e1000532.

105. O’Meara TR, Robbins N, Cowen LE. 2017. The Hsp90 chaperone network modulates *Candida* virulence traits. Trends Microbiol 25:809–819

106. Robbins N, Uppuluri P, Nett J, Rajendran R, Ramage G, Lopez-Ribot JL, Andes D, Cowen LE. 2011. Hsp90 governs dispersion and drug resistance of fungal biofilms. PLoS Pathog 7:e1002257.

107. Li X, Robbins N, O’Meara TR, Cowen LE. 2017. Extensive functional redundancy in the regulation of *Candida albicans* drug resistance and morphogenesis by lysine deacetylases Hos2, Hda1, Rpd3 and Rpd31. Mol Microbiol 103:635–656.

108. LaFayette SL, Collins C, Zaas AK, Schell WA, Betancourt-Quiroz M, Gunatilaka AAL, Perfect JR, Cowen LE. 2010. PKC signaling regulates drug resistance of the fungal pathogen *Candida albicans* via circuitry comprised of Mkc1, calcineurin, and Hsp90. PLoS Pathog 6:e1001069.

109. Diezmann S, Michaut M, Shapiro RS, Bader GD, Cowen LE. 2012. Mapping the Hsp90 genetic interaction network in *Candida albicans* reveals environmental contingency and rewired circuitry. PLoS Genet 8:e1002562.

110. Bielska E, May RC. 2019. Extracellular vesicles of human pathogenic fungi. Curr Opin Microbiol 52:90–99.

111. Zamith-Miranda D, Nimrichter L, Rodrigues ML, Nosanchuk JD. 2018. Fungal extracellular vesicles: modulating host–pathogen interactions by both the fungus and the host. Microbes Infect 20:501–504.

112. de Toledo Martins S, Szwarc P, Goldenberg S, Alves LR. 2019. Extracellular vesicles in fungi: composition and functions. Curr Top Microbiol Immunol 422:45–59.

113. Konečná K, Klimentová J, Benada O, Němečková I, Janďourek O, Jílek P, Vejsová M. 2019. A comparative analysis of protein virulence factors released via extracellular vesicles in two *Candida albicans* strains cultivated in a nutrient-limited medium. Microb Pathog 136:103666.

114. Bielska E, Sisquella MA, Aldeieg M, Birch C, O’Donoghue EJ, May RC. 2018. Pathogen-derived extracellular vesicles mediate virulence in the fatal human pathogen *Cryptococcus gattii*. Nat Commun 9:1556.

115. Freitas MS, Bonato VLD, Pessoni AM, Rodrigues ML, Casadevall A, Almeida F. 2019. Fungal extracellular vesicles as potential targets for immune interventions. mSphere 4:e00747–19

116. Joffe LS, Nimrichter L, Rodrigues ML, Del Poeta M. 2016. Potential roles of fungal extracellular vesicles during infection. mSphere 1:e00099–16

117. Vargas G, Rocha JDB, Oliveira DL, Albuquerque PC, Frases S, Santos SS, Nosanchuk JD, Gomes AMO, Medeiros LCAS, Miranda K, Sobreira TJP, Nakayasu ES, Arigi EA, Casadevall A, Guimaraes AJ, Rodrigues ML, Freire-de-Lima CG, Almeida IC, Nimrichter L. 2015. Compositional and immunobiological analyses of extracellular vesicles released by *Candida albicans*. Cell Microbiol 17:389–407.

118. Oliveira DL, Freire-de-Lima CG, Nosanchuk JD, Casadevall A, Rodrigues ML, Nimrichter L. 2010. Extracellular vesicles from *Cryptococcus neoformans* modulate macrophage functions. Infect Immun 78:1601–1609.

119. Rodrigues ML, Nimrichter L, Oliveira DL, Nosanchuk JD, Casadevall A. 2008. Vesicular trans-cell wall transport in fungi: a mechanism for the delivery of virulence-associated macromolecules? Lipid Insights 2:27–40.

120. Vallejo MC, Matsuo AL, Ganiko L, Medeiros LCS, Miranda K, Silva LS, Freymüller-Haapalainen E, Sinigaglia-Coimbra R, Almeida IC, Puccia R. 2011. The pathogenic fungus *Paracoccidioides brasiliensis* exports extracellular vesicles containing highly immunogenic α-Galactosyl epitopes. Eukaryot Cell 10:343–351.

121. Zarnowski R, Sanchez H, Covelli AS, Dominguez E, Jaromin A, Berhardt J, Heiss C, Azadi P, Mitchell A, Andes DR. 2018. *Candida albicans* biofilm–induced vesicles confer drug resistance through matrix biogenesis. PLoS Biol 16:e2006872.

122. Rodrigues ML, Nimrichter L, Oliveira DL, Frases S, Miranda K, Zaragoza O, Alvarez M, Nakouzi A, Feldmesser M, Casadevall A. 2007. Vesicular polysaccharide export in *Cryptococcus neoformans* is a eukaryotic solution to the problem of fungal trans-cell wall transport. Eukaryot Cell 6:48–59.

123. Rodrigues ML, Nakayasu ES, Oliveira DL, Nimrichter L, Nosanchuk JD, Almeida IC, Casadevall A. 2008. Extracellular vesicles produced by *Cryptococcus neoformans* contain protein components associated with virulence. Eukaryot Cell 7:58–67.

124. Sharov G, Voltz K, Durand A, Kolesnikova O, Papai G, Myasnikov AG, Dejaegere A, Ben Shem A, Schultz P. 2017. Structure of the transcription activator target Tra1 within the chromatin modifying complex SAGA. Nat Commun 8:1556.

125. Peng D, Tarleton R. 2015. EuPaGDT: a web tool tailored to design CRISPR guide RNAs for eukaryotic pathogens. Microb Genom 1:e000033.

126. Shapiro RS, Chavez A, Porter CBM, Hamblin M, Kaas CS, DiCarlo JE, Zeng G, Xu X, Revtovich AV, Kirienko NV, Wang Y, Church GM, Collins JJ. 2018. A CRISPR-Cas9-based gene drive platform for genetic interaction analysis in *Candida albicans*. Nat Microbiol 3:73– 82.

127. Gibson DG, Young L, Chuang R-Y, Venter JC, Hutchison CA 3rd, Smith HO. 2009. Enzymatic assembly of DNA molecules up to several hundred kilobases. Nat Methods 6:343–345.

128. Engler C, Kandzia R, Marillonnet S. 2008. A one pot, one step, precision cloning method with high throughput capability. PLoS One 3:e3647.

129. Edgar R, Domrachev M, Lash AE. 2002. Gene Expression Omnibus: NCBI gene expression and hybridization array data repository. Nucleic Acids Res 30:207–210.

130. Bolger AM, Lohse M, Usadel B. 2014. Trimmomatic: a flexible trimmer for Illumina sequence data. Bioinformatics 30:2114–2120.

131. Dobin A, Davis CA, Schlesinger F, Drenkow J, Zaleski C, Jha S, Batut P, Chaisson M, Gingeras TR. 2013. STAR: ultrafast universal RNA-seq aligner. Bioinformatics 29:15–21.

132. Liao Y, Smyth GK, Shi W. 2014. featureCounts: an efficient general purpose program for assigning sequence reads to genomic features. Bioinformatics 30:923–930.

133. Love MI, Huber W, Anders S. 2014. Moderated estimation of fold change and dispersion for RNA-seq data with DESeq2. Genome Biol 15:550.

134. Goedhart J, Luijsterburg MS. 2020. VolcaNoseR is a web app for creating, exploring, labeling and sharing volcano plots. Sci Rep 10:20560.

